# Breaking Lander-Waterman’s Coverage Bound

**DOI:** 10.1101/060384

**Authors:** D. Nashtaali, S.A. Motahari, B.H. Khalaj

## Abstract

Lander-Waterman’s coverage bound establishes the total number of reads required to cover the whole genome of size *G* bases. In fact, their bound is a direct consequence of the well-known solution to the coupon collector’s problem which proves that for such genome, the total number of bases to be sequenced should be *O* (*G* ln *G*). Although the result leads to a tight bound, it is based on a tacit assumption that the set of reads are first collected through a sequencing process and then are processed through a computation process, i.e., there are two different machines: one for sequencing and one for processing. In this paper, we present a significant improvement compared to Lander-Waterman’s result and prove that by combining the sequencing and computing processes, one can re-sequence the whole genome with as low as *O*(*G*) sequenced bases in total. Our approach also dramatically reduces the required computational power for the combined process. Simulation results are performed on real genomes with different sequencing error rates. The results support our theory predicting the log *G* improvement on coverage bound and corresponding reduction in the total number of bases required to be sequenced.

## I. INTRODUCTION

Data generated from DNA sequencing machines are growing at an unprecedented rate. Extracting knowledge from these data is extremely tedious and usually requires very powerful computing machines. The main reason is that the volume of data generated for an experiment usually contains redundant data and one needs to pay the price of extracting useful information and removing redundant information at the processing step. As an example, in the whole genome sequencing of Human genome with 100x coverage, each base averagely is present in 100 sequencing reads which means 99 percent of the data is redundant. The first question that comes in mind is whether the volume of data generated by the sequencing machines can be reduced without affecting the overall performance. In this paper, we focus on the whole genome sequencing problem and seek fundamental results on the redundancy level required to obtain the desired result.

The first fundamental result in this area has been due to Lander and Waterman in [1] where they present a lower bound on the number of reads, *N*, required to assemble the whole genome. We refer to this bound as *coverage bound*. The coverage bound states that for a genome of size *G* and reads of length *L*, at least 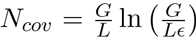 reads are needed such that the whole genome is covered with a probability of no less than 1 – *ϵ* [2]. Therefore, we should have *N* ≥ *N_cov_*.

For the aforementioned scenario, the total number of bases sequenced by the sequencing machine is *NL*, which requires to be of the order of *G* log *G*, from Lander-Waterman’s result. Consequently, the non-reducible redundancy level in such setup will be of the order of log *G*. However, such result is based on the underlying assumption that sequencing and computing steps are performed independently, i.e., a machine takes samples from the genome and sequences as many reads required to cover the whole genome and then another machine processes the reads to assemble the genome. The overall architecture of such approach for whole genome sequencing is shown in Figure 1 (a) and consists of the cascade of two blocks one for sampling and sequencing and one for assembly.

Although such separation between sequencing and processing has been traditionally assumed in the literature, one can question whether such separation is fundamentally optimal with respect to amount of sequenced data which is generated and subsequently processed or not. In other words, can we improve the overall performance of such system by merging the two components? In fact, in order to verify the level of performance improvement achieved by such integration, one needs to answer the following two key questions. First, is there any improvement on lowering the redundancy of generated data by merging the two functions? Second, is it physically possible to build such a machine to perform both functions simultaneously? In the rest of this paper, we will try to answer the first question by proving that one can break the coverage bound of Lander and Waterman and reduce the number of sequenced bases to as low as *O*(*G*). It is worth mentioning that even if we may not have access to a machine that can efficiently combine the sampling and computing units, the approach proposed in this paper can still reduce the computational power required to assemble the genome. This is due to the fact that the processing unit processes the data on the fly and if it detects that the information from the remaining part of data is redundant it will stop further processing of that piece of data. As we will show, such early termination can have significant effect on reducing computational complexity of the whole process. Answering the second question is in fact beyond the scope of this paper. However, our approach clearly shows that if a sequencing machine can be built that can preempt sequencing at the instant the computation part sees best fit, a much more efficient sequencing machine may actually be obtained.

**Fig. 1.**
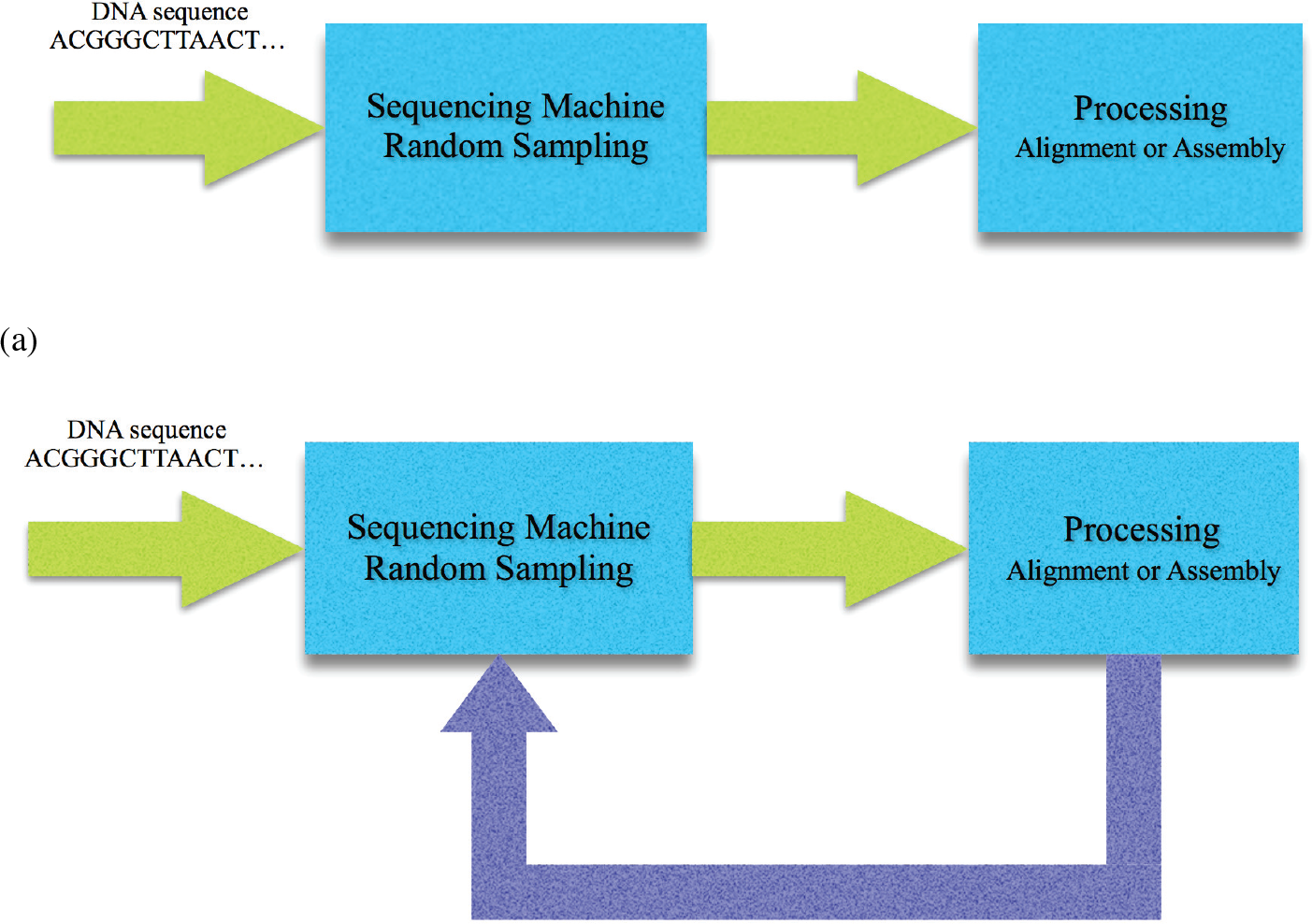
Whole genome shotgun sequencing and our method platforms. In the classic method, the processing starts after termination of random sampling and sequencing. In our proposed sequencing platform, sampling and processing machines work cooperatively. The results of alignment are fed back to preempt reading redundant data in the sequencing machine.

Our strategy to merge the two functions is shown in Figure 1 (b) where the processing machine controls the sequencing machine by blocking sequencing of redundant bases. Hence, we assume that the sequencing machine sequences the DNA fragments base by base and it will stop sequencing a fragment once a blocking command from the processing machine is initiated for that fragment.

In this paper, we only focus on the re-sequencing problem in the processing machine. we first present theoretical results for i.i.d. genome and real genomes with given repeat structures with noiseless reads, and i.i.d. genome with noisy reads. Our simulation results performed on chr19 of Human genome hg19 for both noiseless and noisy reads. We have shown significant improvement on coverage bound for real genome is achieved by using this method. For processing machine, Meta-aligner [3] are used to align reads on the reference genome.

The longer the read lengths, more reduction on number of read bases will be achieved by our method. A number of Next Generation Sequencing (NGS) methods [4], such as PacBio [5] and Nanopore [6]–[8] already provide reads of several thousand bases long and are suitable candidates for such analysis. Figure 2 shows read length distribution of PacBio technology for the first two read archive in NCBI GenBank SRX533609 [9]. It can be envisioned that other sequencing methods might also provide longer reads as sequencing technology further advances in that direction in years to come.

The organization of this paper is as follows. Our basic method for i.i.d. and real genomes with noiseless reads, and i.i.d. genomes with noisy reads are discussed in the Methods section. The simulation results on real genome by considering different sequencing error rates are presented in the Results and discussion section. The last section is devoted to the conclusion and future works.

**Fig. 2.**
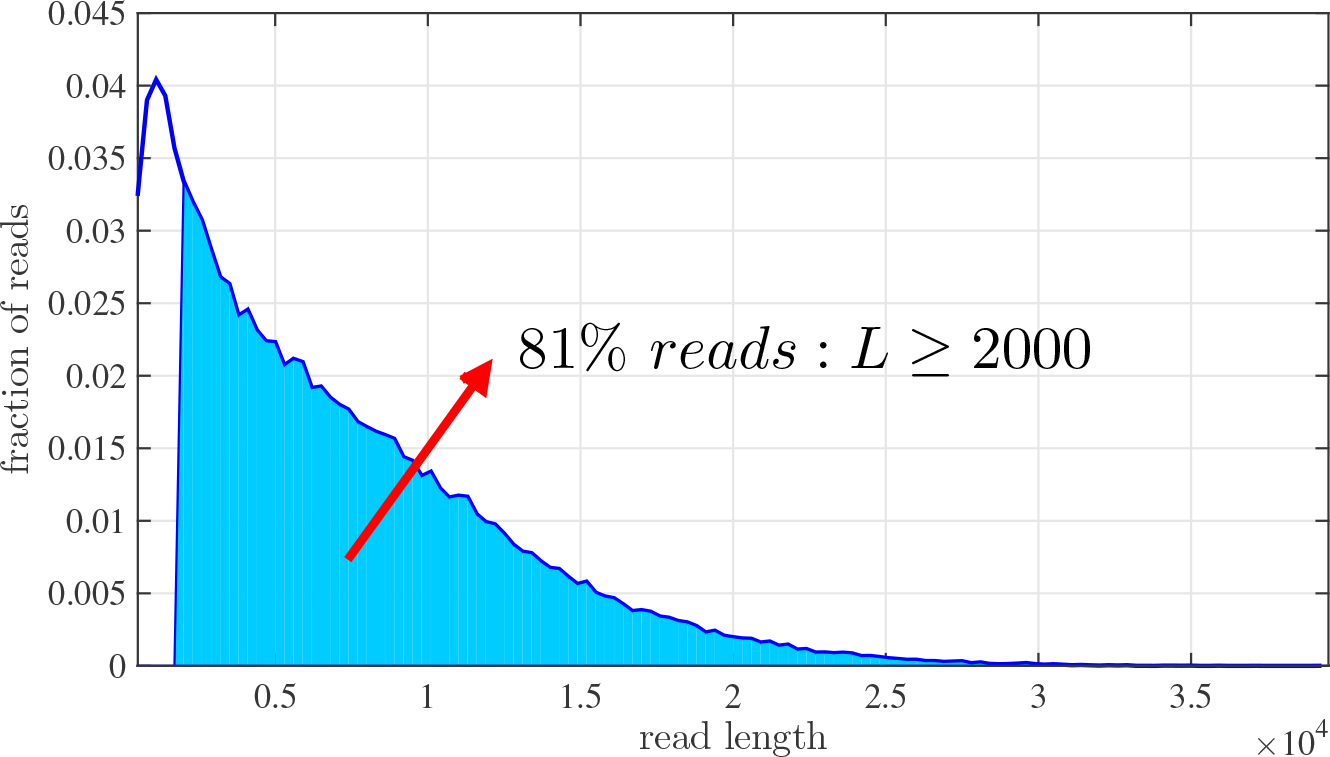
PacBio read length distribution. Almost 81% of PacBio reads have length of at least 2000 bps.

## II. METHODS

Conventionally, in the re-sequencing problem, a reference genome and a set of reads from a target genome are available at the processing step. In this framework, the sequencing machine produces reads of length *L* from *N* DNA fragments. Due to Lander-Waterman’s coverage bound, *NL*, the total number of bases read by the machine, is required to be *O*(*G* log *G*).

Our method changes this bound by assuming that sequencing can be controlled by a processing machine such that it can terminate sequencing at any base. In other words, the sequencer starts reading bases of one side of a DNA fragment one by one and it will stop as soon as a command is initiated by the processing machine. In order to explain the basic ideas and to prove that sequencing can be performed efficiently, We first analyse our proposed methods on i.i.d. genomes. We extend the results to real genomes where repeats play an important role in the structure of the genomes.

### A. I.i.d. genomes

In this part, we assume that the reference and target genomes are i.i.d. random sequences of {*A, C, G, T*} with uniform base probabilities. We also assume that reads are sampled uniformly and independently from the target genome. Consider that reads are noiseless. The key strategy that we use to terminate the sequencing process in a controlled manner is as follows. We divide *L* by some integer number *K* ∈ {0,…, *L*} and without loss of generality assume that *ℓ* = *L/K* is also an integer. We allow the reading machine to read the first *ℓ* bases of all the DNA fragments. Let 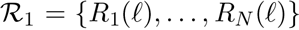 denote the set of starting *ℓ* bases of all the fragments. Here, *R_i_*(*ℓ*) is the first *ℓ* bases of the *i*^th^ fragment.

After generating 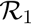, all reads are mapped to the reference genome. Some of the the reads can be mapped uniquely to a location on the genome. We call such reads *anchored*. More precisely, a read *R* is assumed to be anchored if there is only one location on the genome with Hamming distance no more than *α*|*R*| where |*R*| is the read length and *α* is some fixed constant.

After mapping, we partition the set 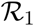 into three disjoint sets: 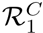 the set of reads that are anchored to some location on the genome and in addition, extending them does not increase the coverage, 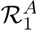 the set of reads anchored to some location on the genome and in addition, extending them will increase the coverage, and 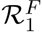 the set of reads that are not anchored in the first step. For a read *R_i_*(*ℓ*) in 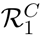 a termination command is initiated to stop further reading of the *i*^th^ fragment. The union of 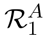 and 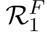 is denoted by 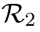 that is the set of fragments where reading processes will be continued on them.

Subsequently, the next base of all fragments in 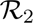 are read and we use the same procedure for mapping and termination. Therefore, at the end of this step, we end up with the set 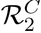 of anchored and terminated fragments with length *ℓ* + 1 and the set 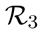 that is used for extension in the next step. In this way, one can proceed to step *L* – *ℓ* + 1 where all the fragments are extended to the maximum length *L*.

If we denote the set of reads that are uniquely mapped in the algorithm by 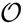, then 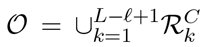. Our proposed algorithm is then detailed in Algorithm 1.

**Algorithm 1.**
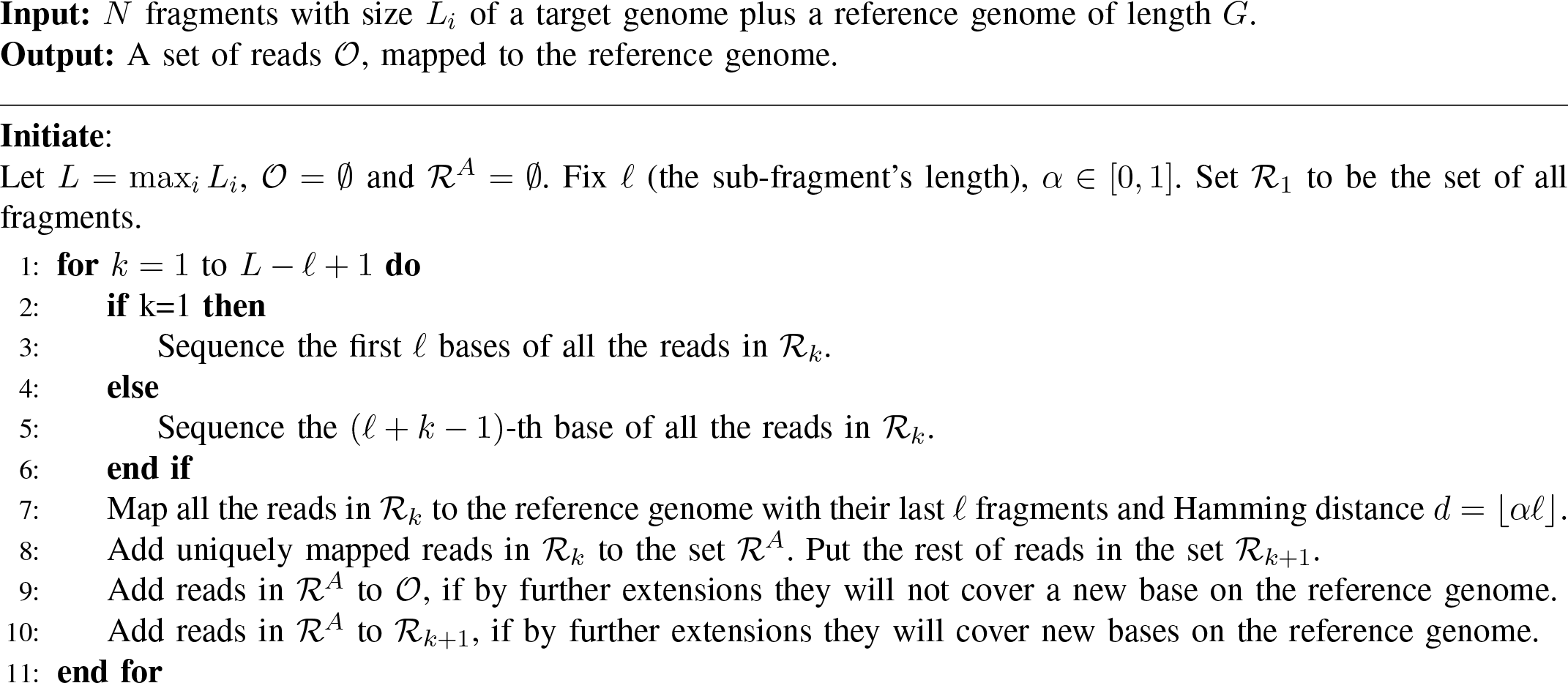

The two parameters of the algorithm, i.e., *ℓ* and α, should be specified based on the structure of reference genome as well as, *G*, *N* and *L*. One can choose *ℓ* to be as low as 1. However, the chance of finding uniquely mapped reads is very small for short reads resulting in much higher processing time. On the other hand, if *ℓ* is chosen to be large, then a lot of reads will overlap after the first step of the algorithm and we will sample a lot of redundant bases. Therefore, an optimal choice of *ℓ* is desired. The optimal value of *ℓ* depends on the size and repeat structure of the genome as well as the statistics of the variations between target and reference genomes. In the following, we describe selection of these parameters for noiseless reads.

It can be shown that for a given random DNA sequence of length *G*, the probability of observing two exact copies of a substring with length *ℓ* is lower than *G*^2^4^−*ℓ*^ [2]. Hence, a noiseless read of length *ℓ* > log *G* from target genome almost certainly can be uniquely mapped to the reference genome. Conversely, a noiseless read of length *ℓ* < log *G* from target genome will be mapped to at least two locations on the reference genome. Therefore, for noiseless reads with no variation between target and reference genomes, we can choose *ℓ* = log *G*.

The choice of α depends on the mapper quality and allowable Hamming distance between reads and the reference genome. If we assume a perfect mapper, the Hamming distance between a read and its true location is *v*|*R*| where *v* is the variations rate between target and reference genomes. Therefore, it suffices to set *α* = *v*. In this scenario, we consider that variation between reference and target genome is negligible.

In order to evaluate the performance of the algorithm, we need to prove that the genome can be completely covered by the reads in 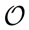 and the number of bases read more than once is small. After the first step of the algorithm, i.e. sequencing *ℓ* bases of all reads, all reads are anchored to their correct location on the genome with maximum Hamming distance of *d* = 0. This is due to the preceding discussion on random genomes where duplicate segments are scarce. We distinguish between two cases:

1. Two subsequent reads have common bases, such as *i*^th^ read and (*i* + 1)^th^ read in Figure 3.
2. Two subsequent reads do not have a common base, such as *j*^th^ read and (*j* + 1)^th^ read in Figure 3.

In the first case, we will encounter reads whose extension does not increase the coverage and therefore, their sequencing should be terminated. These kind of reads belong to 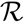. We call the bases common between *i*^th^ and (*i* + 1)^th^ read in such case, over-read bases. In the second case, we will encounter reads whose extension will increase the coverage. We put reads of this kind in 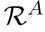, to be further extended in the following steps of the algorithm. Consequently, sequencing of an aligned read in 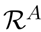 continues until it becomes a member of 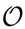.

**Fig. 3.**
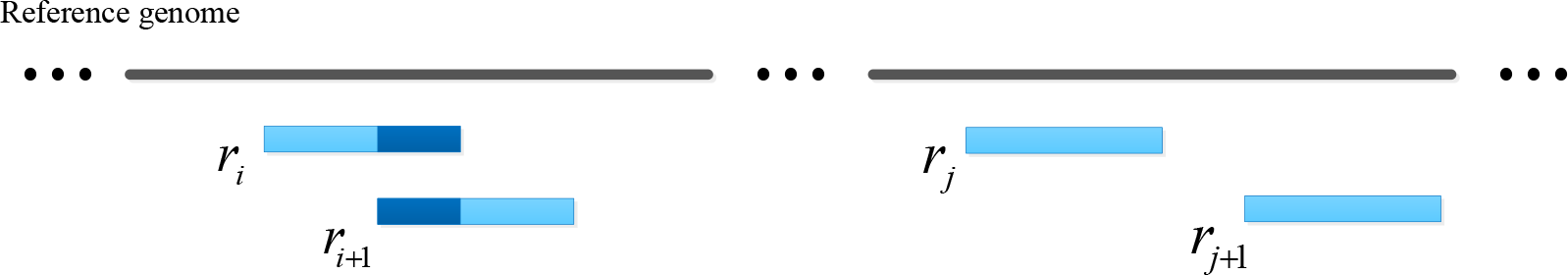
Two possible cases for two subsequent reads. They are either disjoint or have some common bases.

For further analysis, we assume that starting point of reads are a *Poisson* point process with rate 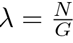. Hence, the inter-arrival times have independent *exponential* distributions.

**Fig. 4.**
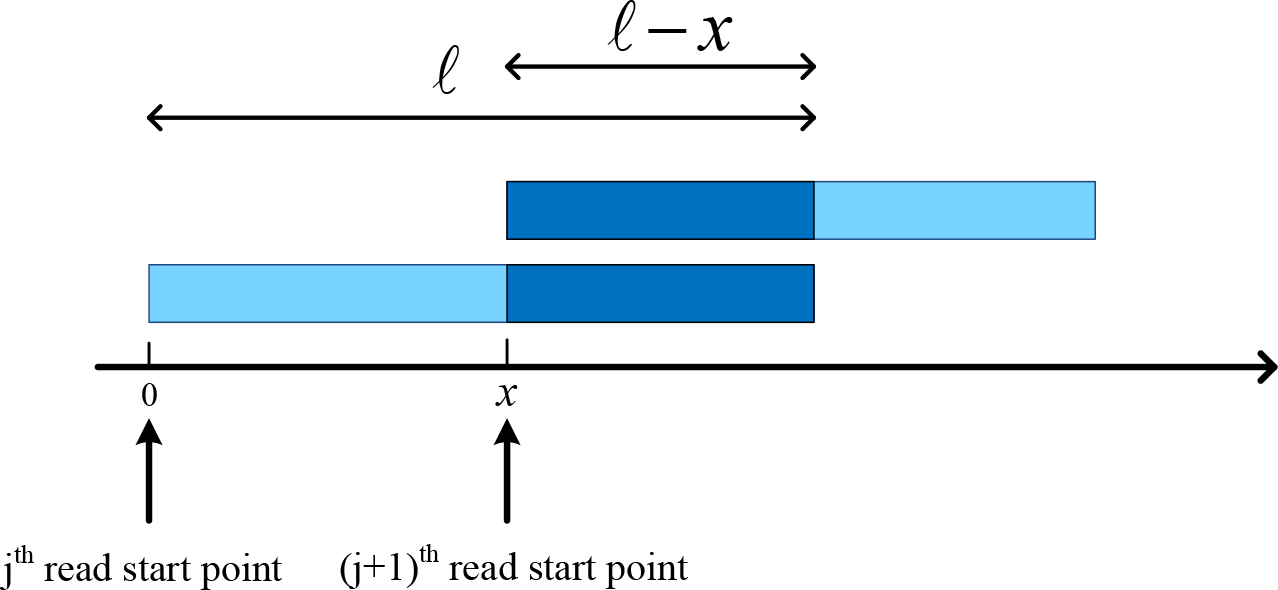
A read of 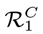 and its adjacent read. The starting point of the second read is *x* bases after the starting point of the overlapped read from the right hand side (assuming that the two reads are of equal length *ℓ*).

Let 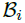 be a random variable representing the number of extra bases read by the sequencing machine for the *i*^th^ read. If 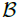 denotes the total number of over-read bases, then 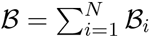. Therefore, 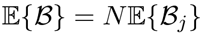.

To compute 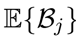, note that the next read of the *j*^th^ read starts *x* bases after the *j*^th^ read. If *x* ≤ *ℓ*, then 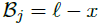. Otherwise, 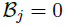, see Figure 4. Since *x* has an exponential distribution, we obtain

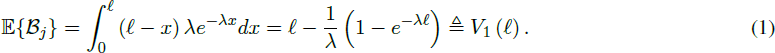

In the case of i.i.d. genomes, we set *ℓ* = log *G*. Hence, the average number of over-read bases of all reads is as follow:

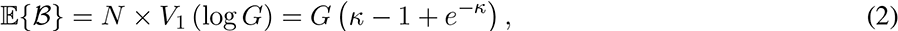

where 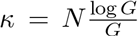. For a constant value *κ*, the order of 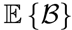 becomes *O*(*G*) and number of reads becomes 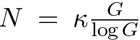. To be able to cover the genome with *N* reads, we need to be able in closing gaps of at least *G* log *G*/*N* bases (from the coverage bound). Therefore, length of reads becomes 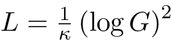.

We observe that by tuning *κ*, one can control maximum read lengths as well as total number of bases read by the machine. For instance, by choosing *κ* = 1, we obtain 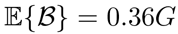 and the maximum read length becomes *L* ≈ 1000 bps.

Next, we consider noisy reads with the error rate *ϵ*. Moreover, we assume that the target and reference genomes vary in their sequences with the variation rate *v*. Before presenting our algorithm for noisy reads, we set a parameter denoted by *C*_*ϵ*_ for the coverage depth. The coverage depth helps in removing sequencing errors by averaging over several reads. The coverage depth is chosen such that if *C*_*ϵ*_ reads cover a base then it is possible to correctly recover that base with a given probability *P*_*ϵ*_.

To compute *C_ϵ_*, we choose a base as the target base if it has maximum vote amongst all other bases over reads covering that location. Let 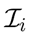 denote the error event of incorrect calling of the *i*^th^ base. If the random variable *C_i_* denotes the coverage of the *i*^th^ base, then error occurs if the corresponding base of more reads are incorrect base. Thus, the probability of error for the *i*^th^ base is,

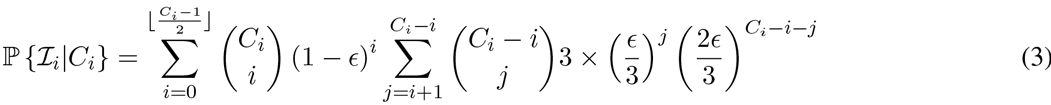

Hence,

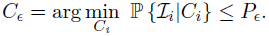

For an example, if *P*_*ϵ*_ = 10^−4^ and *ϵ* = 0.07, *C_ϵ_*becomes 6. For having coverage depth of *C_ϵ_*, it suffices to change the *k*^th^ step of Algorithm 1 as follows: we add a read in to 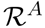 to 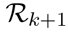 when its extension will cover a base for *c*^th^ times on the reference genome, where *c* ≤ *C_ϵ_*. Details of the proposed algorithm for noisy reads is presented in Algorithm 2.

**Algorithm 2.**
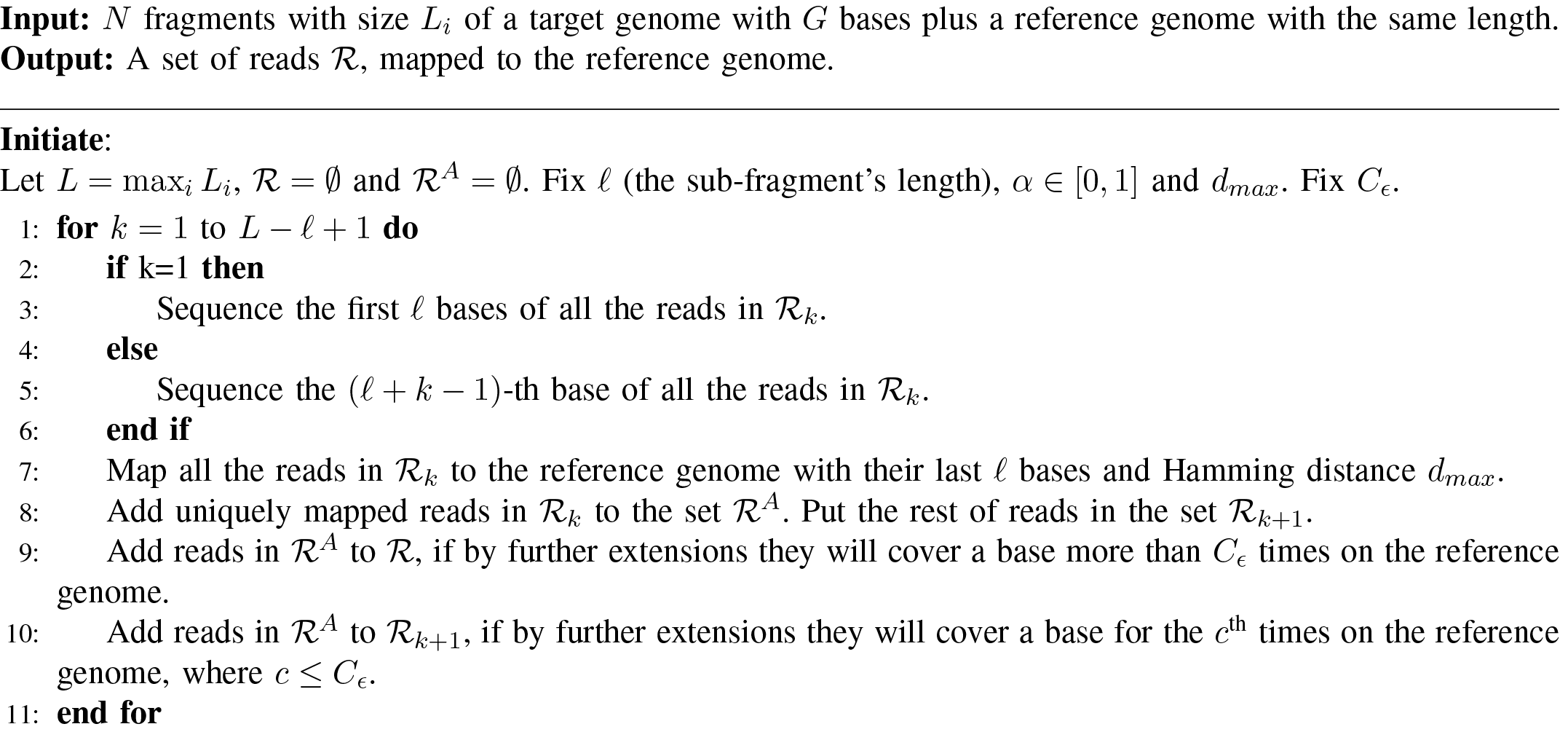

Mapping fragments of length *ℓ* with maximum Hamming distance *d_max_* to the reference genome is error prone and we first analyze the performance of the alignment procedure. Let 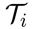 and 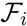 denote the true and false alignment events for the *i*^th^ read, respectively. The *i*^th^ read of length *ℓ* with a maximum Hamming distance of *d_max_* is mapped to its true location with probability

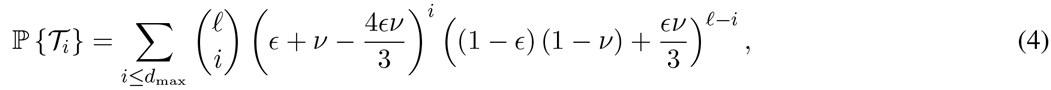

and is mapped to a false location on the genome with maximum Hamming distance *d_max_* with probability (using the *union bound*),

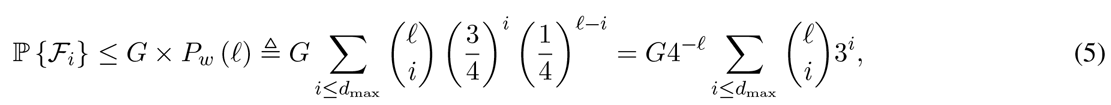

where *p_w_*(*ℓ*) represents the probability of incorrect alignment of a fragment of length *ℓ* to another position on the reference genome. Denote, 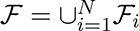 as the false alignment event. Hence,

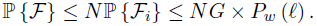

Therefore, if *p_w_*(*ℓ*) scales as 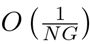,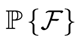 tends to zero and all reads are mapped to their true location with 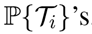. In the worst case, if any read is extracted from each base of the reference genome, i.e. *N* = *G*, then *p_w_*(*ℓ*) scales as 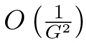. It can be easily verified that for *ϵ* = 0.05, *ℓ* = 2 log *G* and *d_max_* = 8, 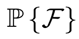 tends to zero and 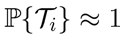. Therefore for *ϵ* ≤ 0.05, all noisy fragments of length *ℓ* are *uniquely* mapped to their correct locations on the reference genome.

To compute the number of extra read bases, we use a similar argument as the one used in the noiseless case. To obtain 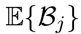 for the *j*^th^ read in this case, assume that the 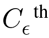 read after this read starts *x* bases further. Again *B_j_* = max{0,*ℓ* – *x*}. Since, *x* has an *Erlang* distribution, we obtain

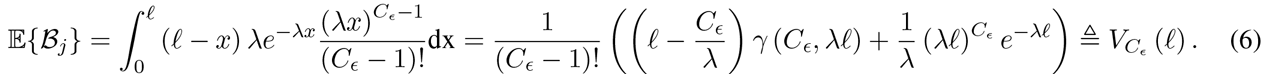

where *γ (s, x)* is the lower incomplete gamma function and for *s* ∈ ℕ is equal to

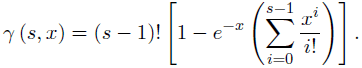

Thus, the average number of over-read bases for all reads in this step is,

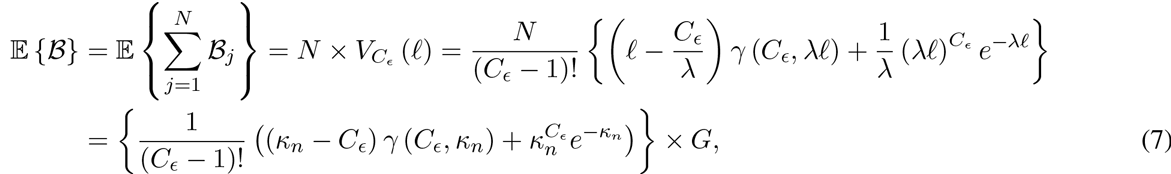

where 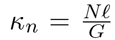. Therefore, for any constant *κ_n_*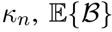 becomes *O*(*G*). The coverage bound when each base of the genome is covered by at least *C_ϵ_* reads can be determined as follows,

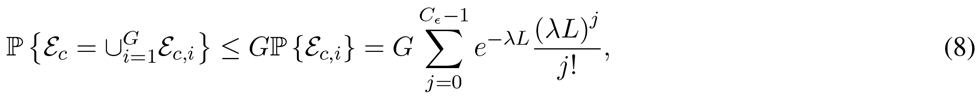

where *ε_c_* and *ε_c,i_* are error events that at least one base and the *i*^th^ base of the genome are not covered by at least *C_ϵ_* reads, respectively. Therefore, if

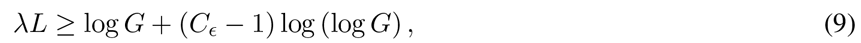

then each base of the genome is covered by at least *C_ϵ_* reads almost surely. Subsequently, for a fix *ℓ* and constant value of *κ_n_*, number of reads and read length from (9) become 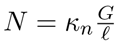 and 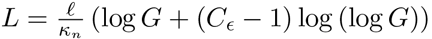 respectively.

Again, there exists a trade-off between the length of reads and number of read bases that can be controlled by *κ_n_*. Figure 5 shows 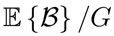 in (7) for sequencing error rates of *ϵ* = {0,0.02,0.05} and *C_ϵ_* = 6. Also, Figure 6 shows total read bases relative to read length for different sequencing error rates *ϵ* = {0,0.02,0.05,0.1}. We set *P*_*ϵ*_ = 10^−4^ and therefore for *ϵ* = {0,0.02,0.05,0.1} we use: *C_ϵ_* = {1, 4,6,8} from (3), and *ℓ* = {1,1.5, 2, 3} ×log *G* with *d_max_* = {0, 4, 8,15} which satisfy alignment constraints, respectively. These figures show that when *ϵ* = 5%, approximately 6.08*G* bases (only 0.08*G* bases are over-read) are read for read length of *L* ≈ 1000 bps.

**Fig. 5.**
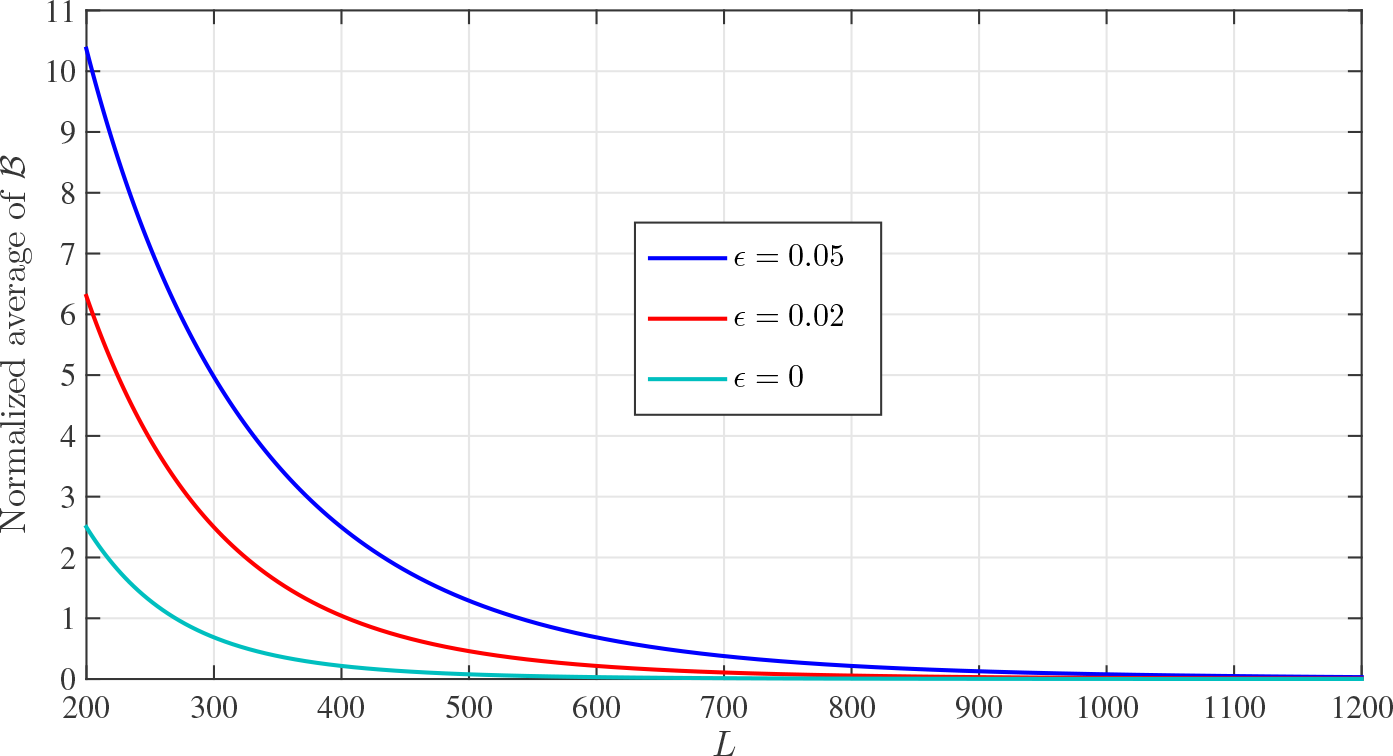
The normalized average number of over-read bases and read lengths for different sequencing error rates and *C_ϵ_* =6.

**Fig. 6.**
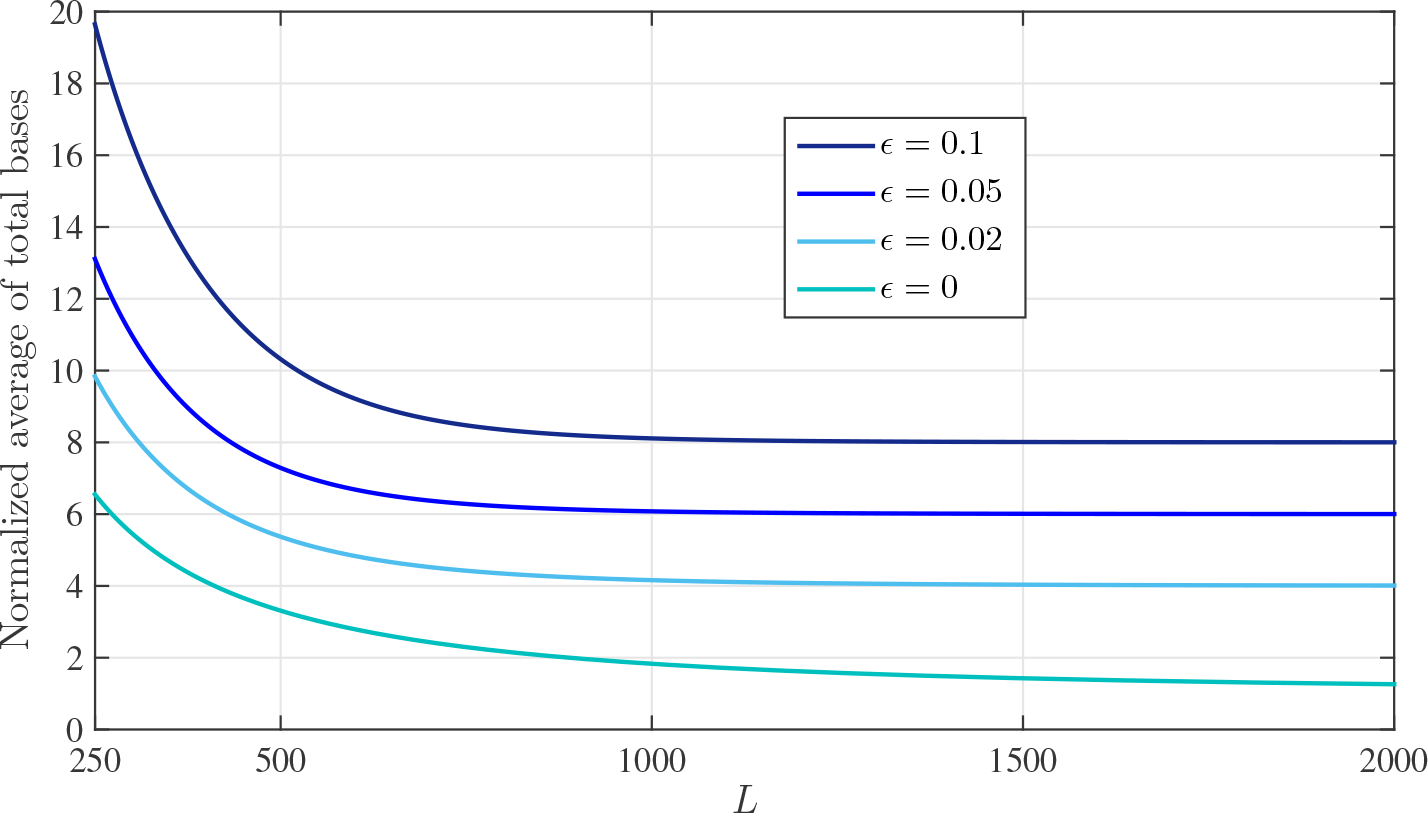
The normalized total number of read bases and read lengths for different sequencing error rates and *P*_*ϵ*_ = 10 ^−4^. Note that, total number of read bases in each error rate tends to its corresponding *C_ϵ_*.

### B. Real Genomes

In this section, we consider DNA sequencing of real genomes where many repeats are dispersed across the genome. First, we assume that reads are noiseless. Note that, if all the *ℓ*-mers of the genome are repetitive elements then Algorithm 1 fails in anchoring reads correctly to the reference genome and therefore reading *O* (*G* log *G*) is unavoidable. However, as we will show, the repeat patterns in real genomes allow successful coverage of bases with only *O* (*G*) reading bases.

A mosaic model for capturing the repeat structure of the genome is presented in [3]. In this model the reference genome consists of two types of intervals, repeat and random intervals. These types are defined based on two parameters *ℓ* and *d*: representing the fragment length and mismatch factor, respectively. Repeat (random) intervals are consecutive bases where any fragment of length *ℓ* starting from a base within these intervals can be aligned to some other location(s) (one location) of the genome with maximum Hamming distance *d*. For the sake of simplicity, we consider only *d* = 0.

Let denote a set of all exact repeat intervals of the reference genome as *ℛ*. Also, assume that a repeat *R* ∈ *ℛ* has length *ℓ_R_* and repeat lengths have the distribution *f_ℓ_*. We need to treat reads starting from repeat and random intervals, differently. For this purpose, we consider three starting regions for start point of a given read, as *S*_1_, *S*_2_, and *S*_3_. We can determine the average number of total over-read bases (i.e. 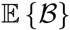) as follows,

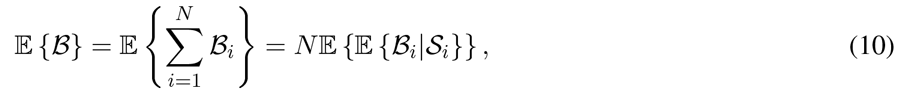

where **S_i_** is a random variable that shows the starting region of the *i*^th^ read. Hence,

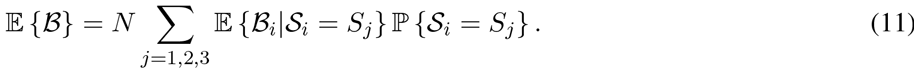

In the following, we determine each term of equation (11). Region *S*_1_ is random intervals such that reads starting from random intervals can be anchored to their true locations based on their first *ℓ* bases. Therefore, we can readily compute the average number of over-read bases in random intervals using equation (1). More precisely,

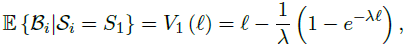

where the total average number of these reads is

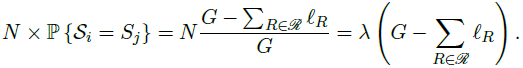

On the other hand, reads starting from repeat intervals can not be anchored unless it contains an *ℓ*-mer which resides in random interval. Using this fact, c.f. Figure 7, each repeat interval of length *ℓ_R_* can be partitioned into two disjoint intervals: 1) Mappable zone: the last min{*L–ℓ, ℓ_R_*} bases, 2) Un-mappable zone: the first max{0, *ℓ_R_–L+ℓ*} bases. Regions *S*2 and *S*3 are mappable and un-mappable zones, respectively.

Clearly, reads from un-mappable zones cannot be anchored and therefore they need to be read up to length *L*. Using (1), we compute the average number of over-read bases for each read in un-mappable zones as:

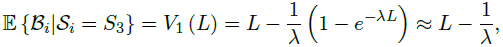

where the total average number of these reads is

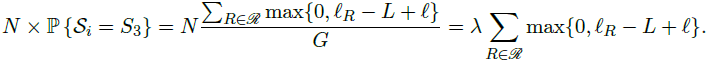

**Fig. 7.**
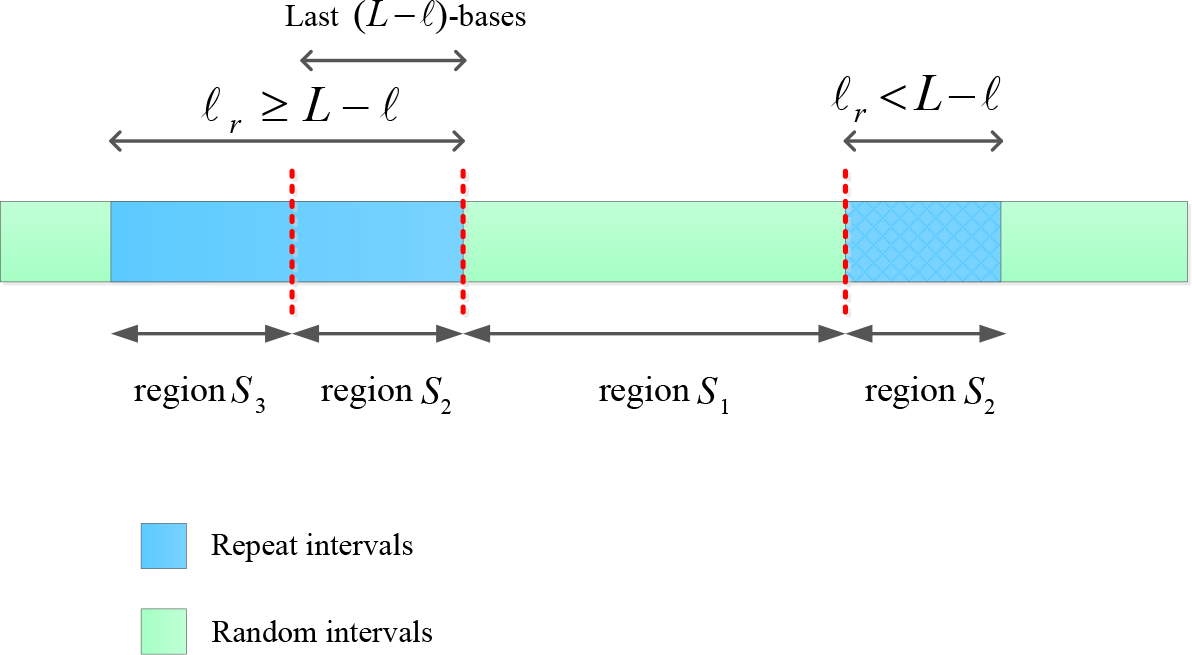
Repeat intervals regions. The first region (*S*_1_) is random intervals. The second region (*S*_2_) is the last *L* – *ℓ* bases of repeat intervals. The third region (*S*_3_) is the other bases of repeat intervals.

It remains to compute the number of over-read bases for reads from mappable zones. Consider all mappable regions *s_m_* ∈ *S*_2_ of length *l_m_*, for *m* = {1,…, |*ℛ*|}. If the *m*^th^ repeat interval has length *ℓ*_*m*_, then = min{*L* – *ℓ,ℓ_m_*}. At least *l_m_* + *ℓ* bases of each read within *s_m_* are being sequenced. Consider the *i*^th^ read has distance of *l_i_* from the end of its mappable zone. The average number of over-read bases for the *i*^th^ read in mappable zones can be determined as:

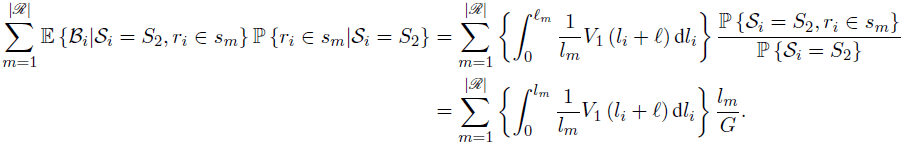

Thus, the average number of total over-read bases for all reads in mappable zones becomes,

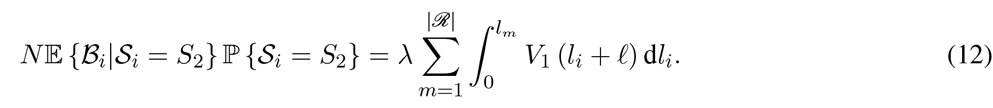

Define,

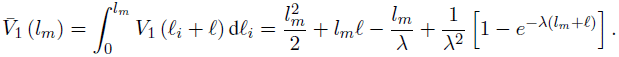

Therefore, the average number of over-read bases by considering repeat structure in (10) becomes,

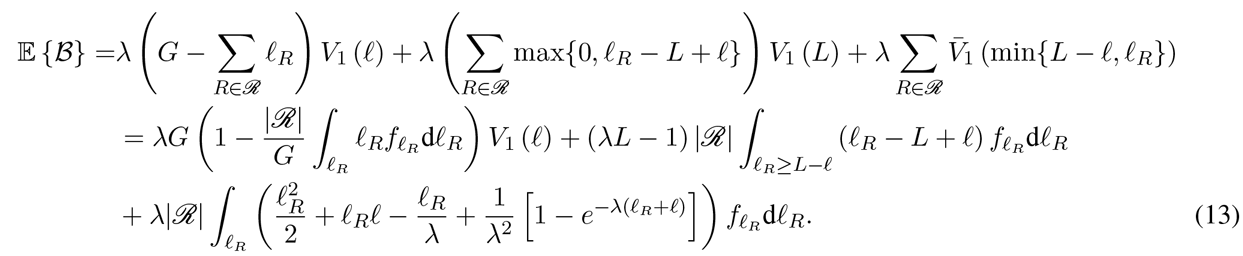

Given the repeat length distribution (i.e. 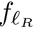) of any real genome, we can determine the 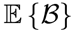 for that genome. The distribution of log 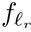 for Human genome hg19 is illustrated in Figure 8. Also, Figure 9 shows 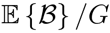 for whole genome of hg19 and i.i.d. genome. In this simulation, we used *ℓ* = log *G* ≈ 30. Results confirm that for real data set, we can read only *O*(*G*) bases to assemble the genome.

**Fig. 8.**
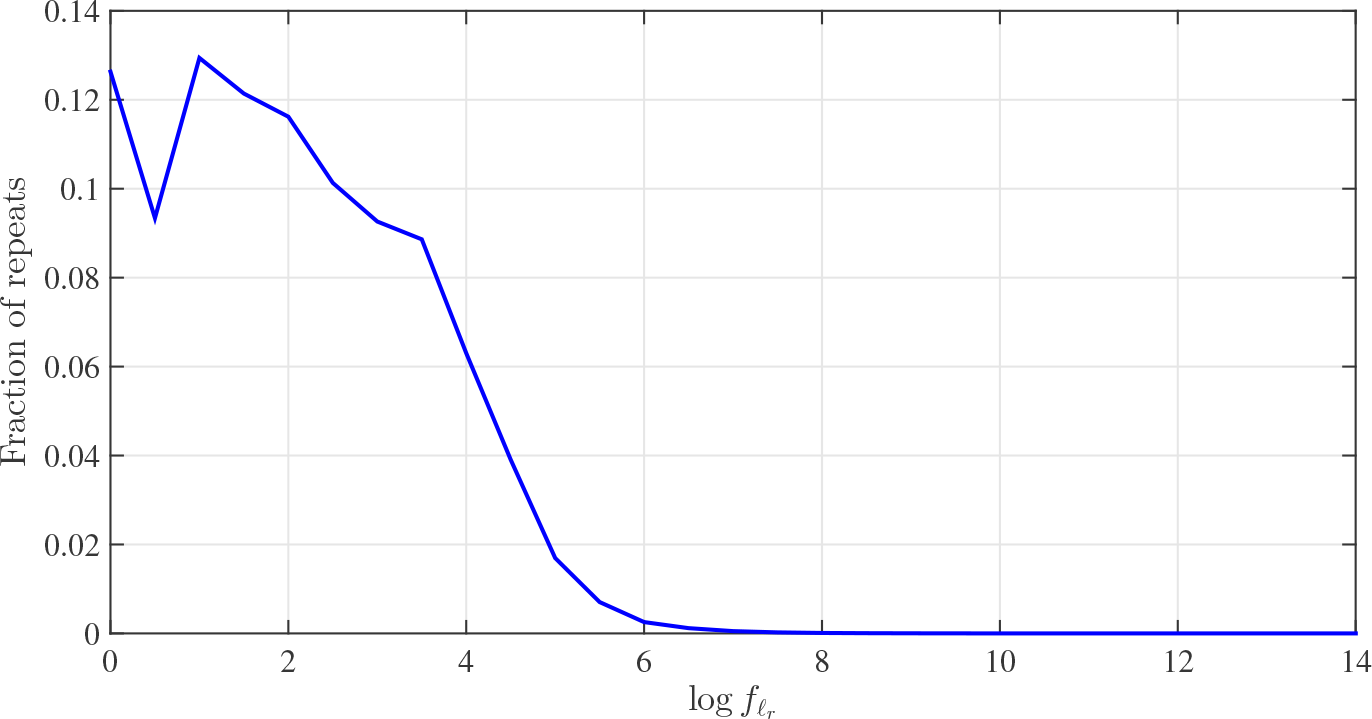
The distribution of repeat lengths logarithm (i.e. log 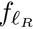) for Human genome hg19 based on Meta-aligner model with (*ℓ, d*) = (30, 0) defined in [3].

When reads are contaminated with sequencing errors of rate *ϵ*, we use a proper value of *ℓ* such that a fragment length of *ℓ* is aligned to its correct location with a probability close to one. Also, consider the coverage depth, i.e. *C_ϵ_*, as (3). We model the reference genome with Meta-aligner (*ℓ, d* = 0)-model. Thus, using the same argument as noiseless reads, the average number of over-read bases can be determined similar to (13) except incorporating 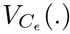 in (6) instead of *V*_1_(.). Therefore, the average number of over-read bases for real genomes in the presence of noise becomes,

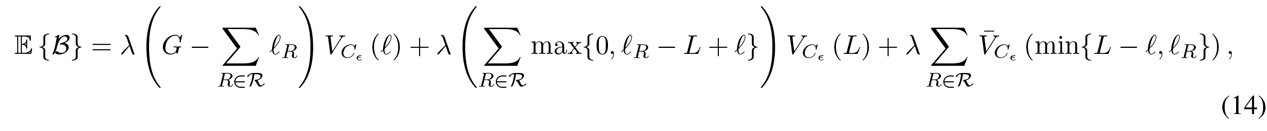

where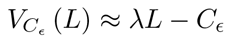 and

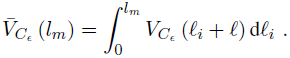

Figure 10 shows the 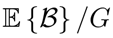 for whole genome of hg19 with *ϵ* = {0,0.05}. We use *ℓ* = log *G* ≈ 30 and *ℓ* = 2 log *G* ≈ 60 for *ϵ* = 0 and *ϵ* = 0.05, respectively.

**Fig. 9.**
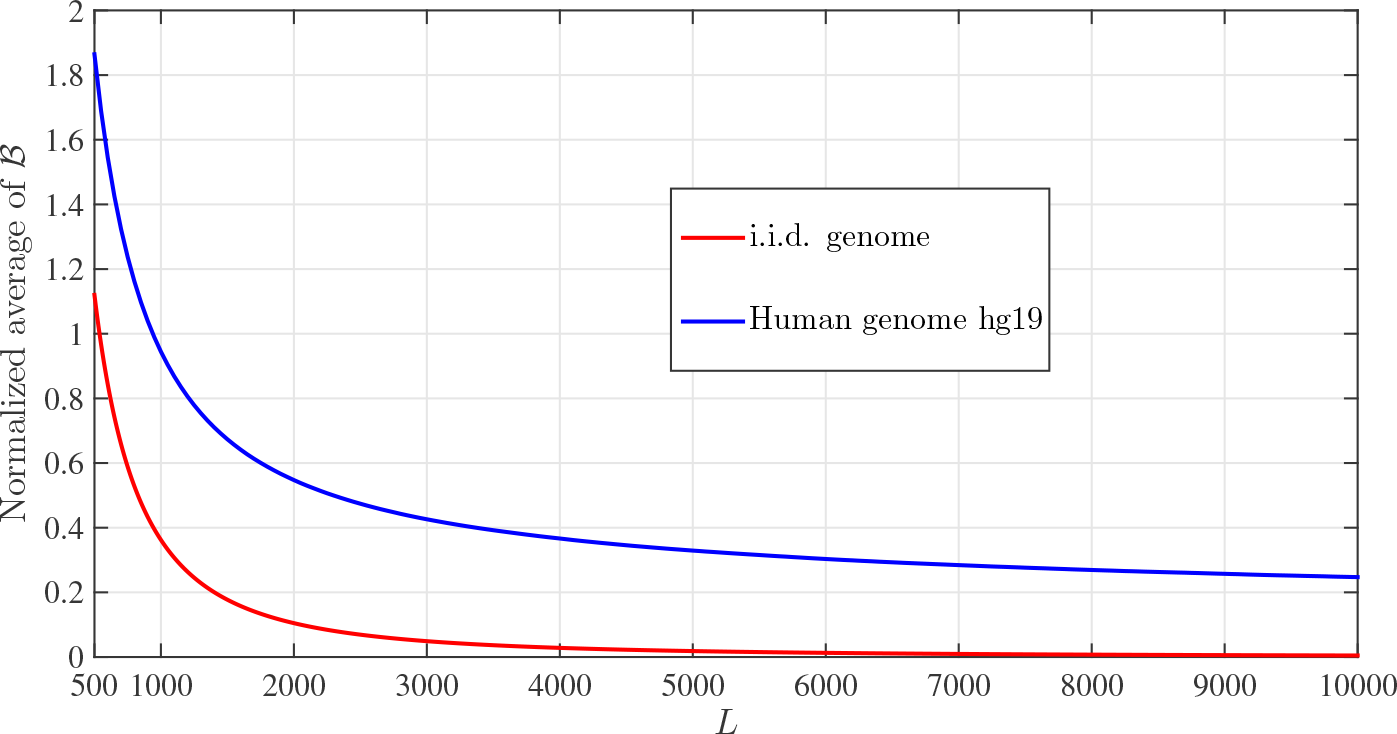
The average number of over-read bases for i.i.d. genome and Human genome hg19 with *ℓ* ≈ 30 and *N* = *G* log *G/L*.

**Fig. 10.**
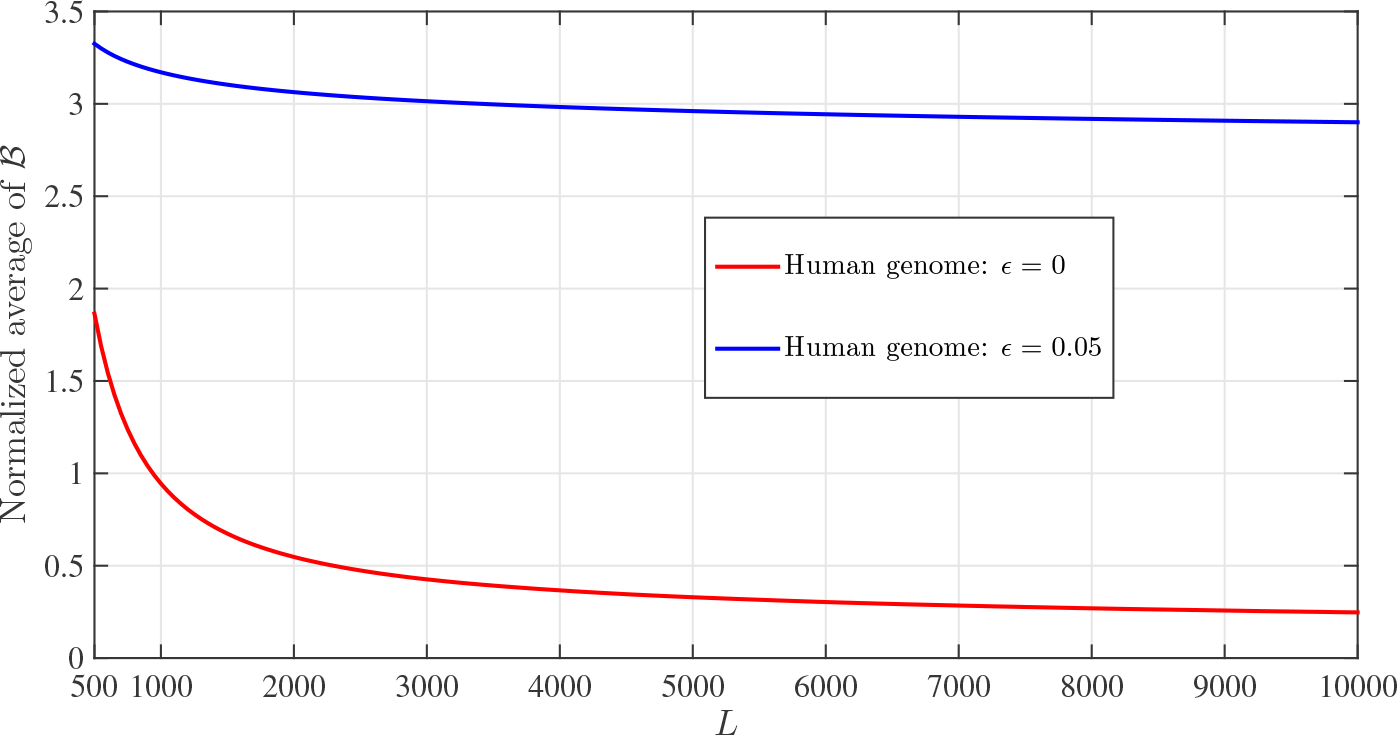
The normalized average number of over-read bases for i.i.d. genome and Human genome hg19 with *ℓ* ≈ {30, 60} and *N* = *G* log *G/L* for *ϵ* = {0, 0.05}, respectively.

## III. ALGORITHM COVERAGE ANALYSIS

In this section, we compute the number of gapped bases in the reference genome when the proposed algorithms are used. First, consider i.i.d. genomes. We show that the whole genome is in fact covered by the noiseless reads when using Algorithm 1. Suppose that all reads of length *ℓ* are aligned to the reference genome. Let *ε* denotes the *error* event which is the event that a base is not covered in our algorithm. Let us denote the event of not covering the *i*^th^ base by *ε_i_*. Thus,

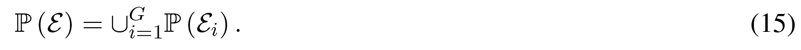

From union bound, we have

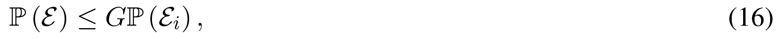

for arbitrary *i* ∈ {1,‥,*G*}. Define the set **S_i_**, consisting of starting points of reads that are aligned to the reference genome with less than *L* bases before the *i*^th^ base location. Since the nearest read in *S_i_* to the *i*^th^ base does not overlap with other reads from its right hand side, this read will be extended in subsequent steps of the algorithm and at the end, the *i*^th^ base will be covered by this read. Hence, *S_i_* must be an empty set and *ε_i_* occurs when no read’s starting point is located less than *L* bases before the *i*^th^ base. This condition is the same as the coverage bound condition. Thus,

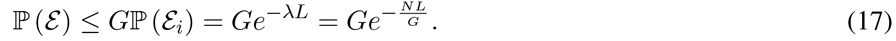

Thus, if the number of reads *N* and the reads’ length *L* satisfy the coverage condition in [1] (i.e. *NL* ≥ *G* log *G*), the sequence is completely covered by the reads in our method for noiseless reads and i.i.d. genomes.

For noisy reads, we show the perfect coverage of reference genome when noisy reads are used in Algorithm 2. For this purpose, the same argument as noiseless reads is considered. Let *ε_n_* denotes the *error* event which is the event that a base is not covered by at least *C_ϵ_* reads in our algorithm. Also, denote error event for the *i*^th^ base by *ε_n,i_*,. Therefore, 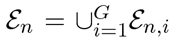 and *ε_n,i_*, occurs when less than *C_ϵ_* read’s starting points are located within *L* bases before the *i*^th^ base. This condition is the same as the coverage bound condition with a given *C_ϵ_* in (8). Thus, if number of reads *N* and reads’ length *L* satisfy the coverage condition for noisy reads in (9), the sequence is completely covered by at least *C_ϵ_* noisy reads in our method as well.

Now consider a real genome. We must determine how many bases are covered with Algorithm 1 or Algorithm 2 (based on noiseless or noisy reads). For this purpose, let *ε_k_* denotes the error event that the *k*^th^ base is not covered by reads with the proposed algorithms. We only consider coverage in this section, therefore, we use *C_ϵ_* = 1 for noisy reads. Based on the base location and its neighboring repeat intervals within the genome, this base is classified to two different classes as shown in Figure 11. Using these two classes, different sub-classes for locating random and repeat intervals can be modelled. Note that similar to i.i.d. genome, locating one read within distance of *L* bases before a given base is sufficient for covering that base. In the following, we determine probability of the *ε_k_* for each class. In the proposed analysis, we assume that each fragment of length *ℓ* is mapped uniquely to the reference genome with probability *p_t_*. Also, we denote the number of reads within a random interval of length *l* as 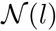.

1. Class A: Assume *d_1_* and *d_2_* bases within distance of *L* bases before and after the *k*^th^ base are in random interval, respectively. If a read has a fragment within a random interval, it can be mapped to the reference genome uniquely. If *d_1_* + *d_2_* ≥ *L*, we divide the interval of length *L* before the *k*^th^ base to three parts: 1) all bases with distance [*L, d_1_*] from base *k*, 2) all bases with distance [*d*_1_, *L* – *d*_2_] from base *k*, and 3) the remaining bases of the random interval with distance [*L – d*_2_,0] from base *k*. Divide the first part to 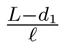 disjoint sub-intervals of length *ℓ*, If any read starts within the *j*^th^ sub-interval, the number of fragments of that read within the random interval is 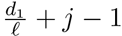. If any read starts within the second part, the number of fragments of that read within the random interval is 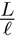. In addition, divide the third part to 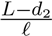 sub-intervals of length *ℓ* such that if any read starts within the *j*^th^ sub-interval, the number of fragments of that read within the random interval is 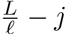. Thus,

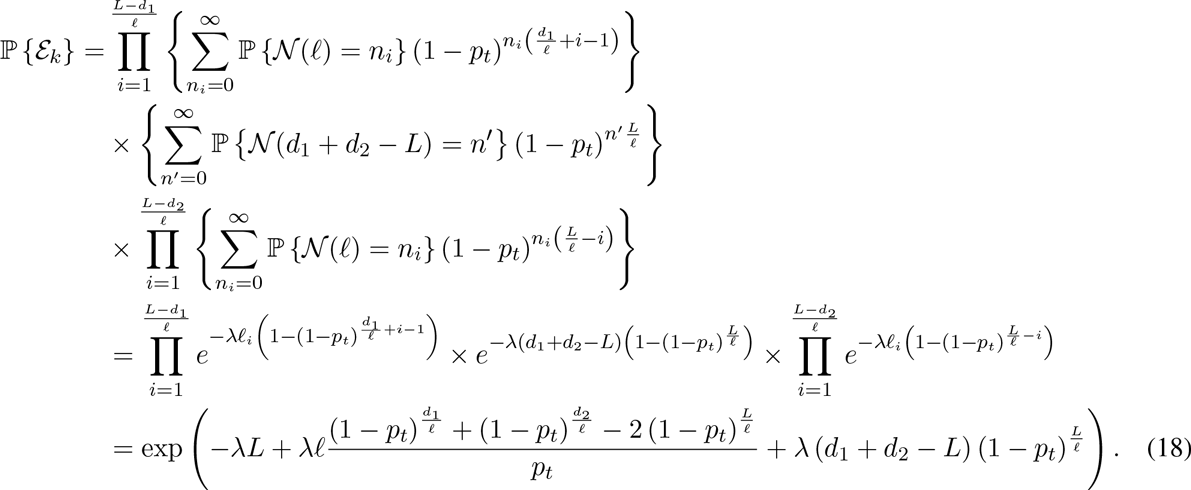 The same result is obtained when *d*_1_ + *d*_2_ < *L*, i.e.,

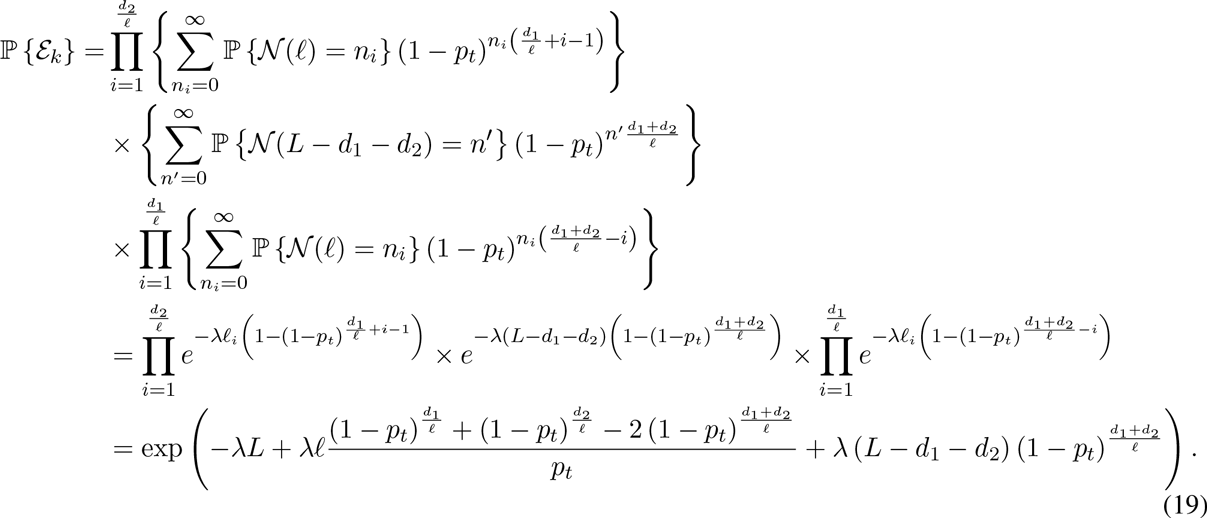
2. Class B: Assume *d*_1_ and *d*_2_ bases within distance of *L* bases before and after the *k*^th^ base are in repeat interval, respectively. If *d*_1_ + *d*_2_ ≥ L, we divide the interval of length *L* before the *k*^th^ base to three parts similar to class A. Thus,

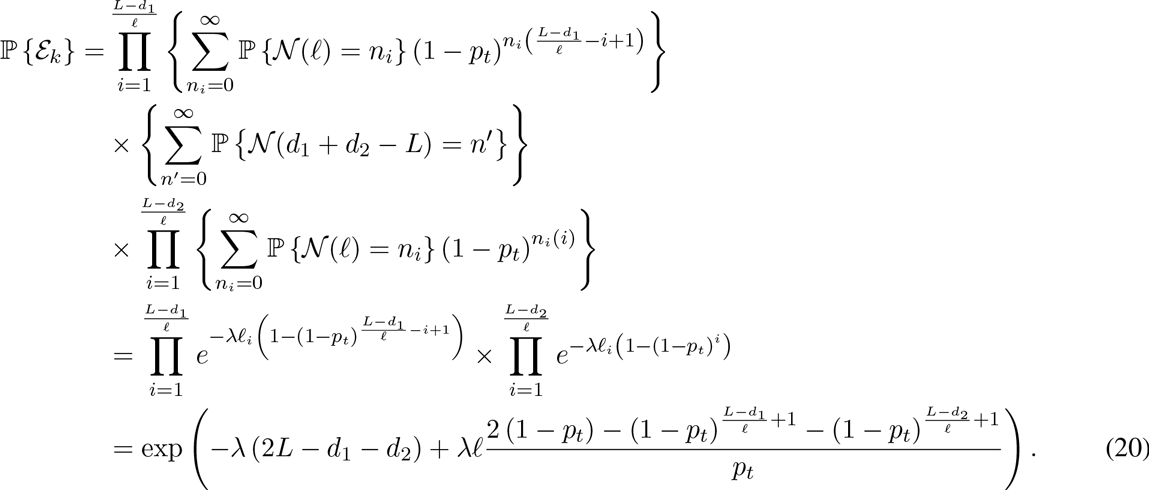

The same result is obtained when *d*_1_ + *d*_2_ < *L*, i.e.,

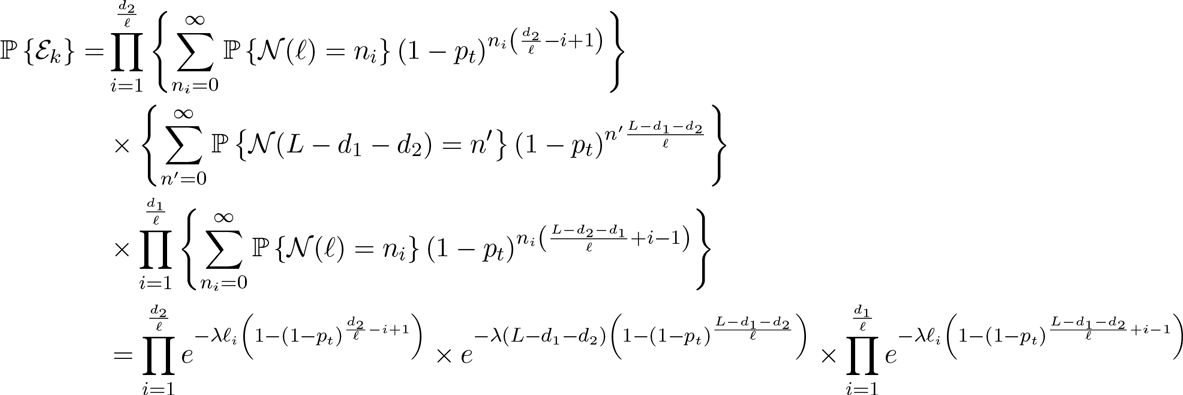

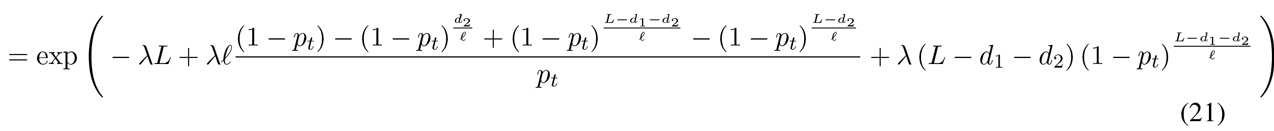

**Fig. 11.**
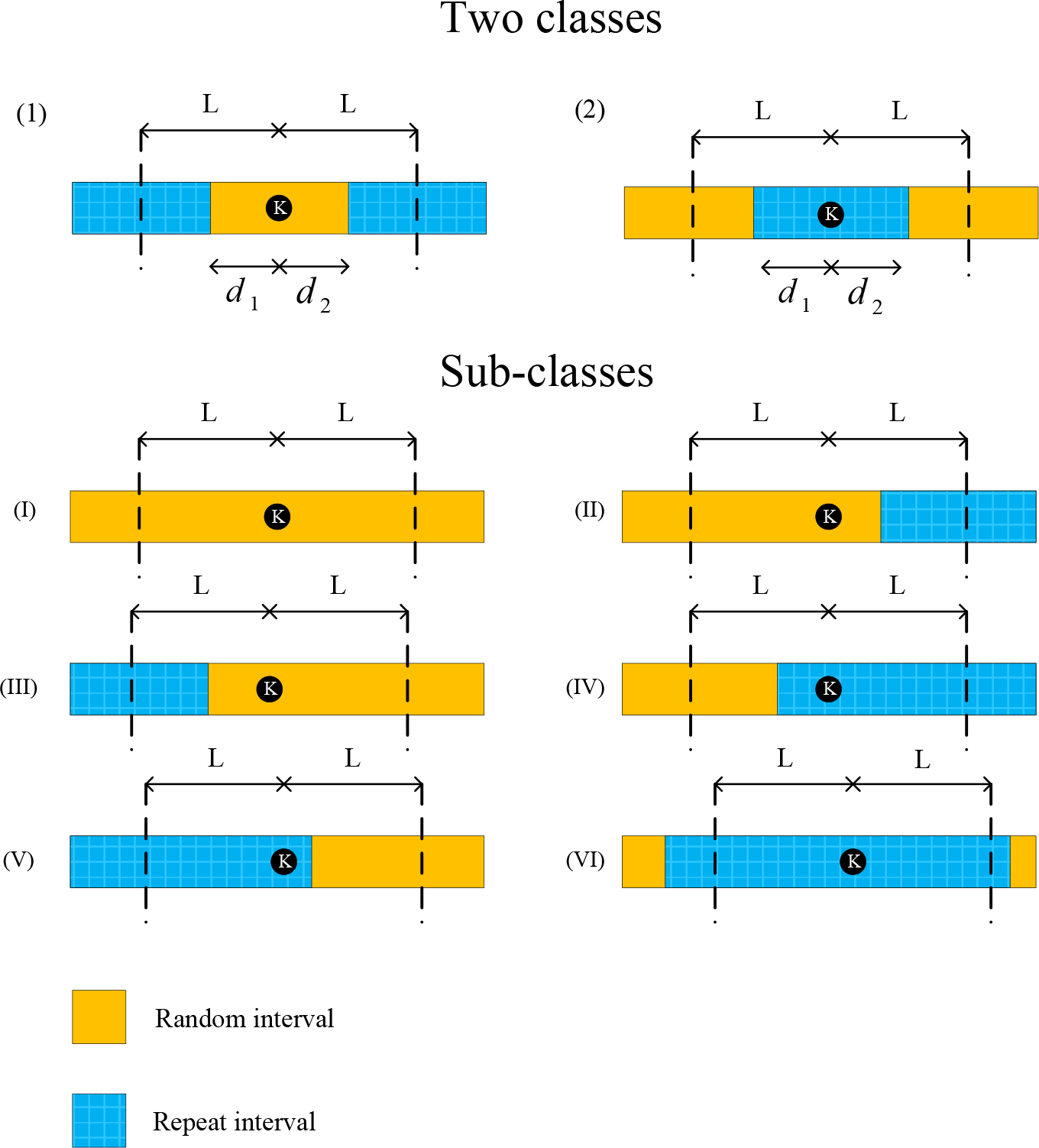
Classification of each base of the reference genome based on random and repeat intervals of the genome. By considering special cases for these two classes, six sub-classes created.

We are interested for error probabilities of some special cases. These interested cases are illustrated as sub-classes in Figure 11. The error probability of each sub-class is determined in the following.

- The first sub-class (I): This sub-class can be modelled with the first class with *d*_1_ = *d*_2_ = *L*. Thus,

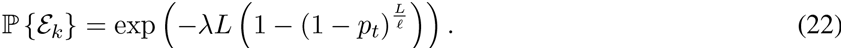
- The second sub-class (II): Consider that *d* bases within distance of *L* bases after the *k*^th^ base is in random interval. This sub-class can be modelled with the first class with *d*_1_ = *L* and *d* = *d*_2_ < *L*. Thus,

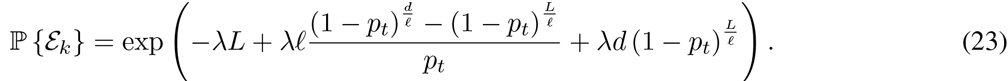
- The third sub-class (III): Consider that *d* bases within distance of *L* bases before the *k*^th^ base is in random interval. This sub-class can be modelled with the first class with *d*_2_ = *L* and *d* = *d*_1_ < *L*. Thus, the error probability of this class is the same as the sub-class (II) except that *L* – *d* bases within distance of *L* before the *k*^th^ base exist in random interval.
- The forth sub-class (IV): Consider that *d* bases within distance of *L* bases before the *k*^th^ base is in repeat interval. This sub-class can be modelled with the second class with *d*_2_ = *L* and *d* = *d*_1_ < *L*. Thus,

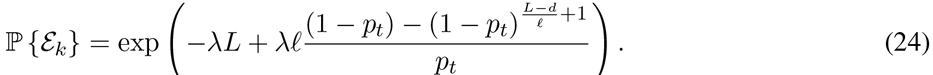
- The fifth sub-class (V): Consider that *d* bases within distance of *L* bases after the *k*^th^ base is in repeat interval. This sub-class can be modelled with the second class with *d*_1_ = *L* and *d* = *d*_2_ < *L*. Thus, the error probability of this class is the same as the sub-class (IV) except that *L* – *d* bases exist within distance of *L* before the *k*^th^ base in repeat interval.
- The sixth class (VI): This sub-class can be modelled with the second class with *d*_1_ = *d*_2_ = *L*. Thus, the error probability of this sub-class is 1.

Thus, by considering repeat structure of the genome, we can determine the probability of coverage for the genome. We classify bases of Human genome hg19 and determine probability of gap for each class using (18)–(21). The average probabilities of gap, i.e. 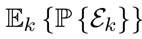, for different values of 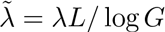 and read lengths (*L*) are shown in Figures 12–13. These probabilities of gap are shown for *p_t_* = 1 and *p_t_* = 0.7 (dotted line). Note that, the coverage bound shows that using reads of length *L* ≥ log *G* and 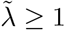, all bases of an i.i.d. genome are covered reads with probability almost one.

## IV. RESULTS AND DISCUSSION

In this section, we propose simulation results for chr19 of Human genome hg19 with different sequencing error rates.

### A. Benchmark

The chr19 of Human genome hg19 is used as the reference. Noisy reads are also extracted uniformly from the reference genome and are mapped to this genome. We consider sequencing error rates of *ϵ* = {0, 5,10}% with 90% mismatches and 10% indels. Errors are added by an i.i.d. manner. We use Meta-aligner [3] to align reads with their two fragments of length *ℓ* not with all their bases. We consider only the first stage Meta-aligner. Since with a mismatch percentage of α and read length of *ℓ*, there are *α* × *ℓ* bases altered in each read on the average, we allow Meta-aligner to align reads to the reference genome with a distance of [*α* × *ℓ*]. For chr19 of Human genome hg19 of size *G* bases, *ℓ* = log *G* is approximately equal to 26 and we use *ℓ* = 30. Therefore, *N* = (30 + 4 (*C_ϵ_* – 1)) *G/L* reads are randomly generated from chr19 for any read length *L* and *C_ϵ_*. Also, we consider *C*_0_ = 1, *C*_5_ = 6, *C*_10_ = 8.

**Fig. 12.**
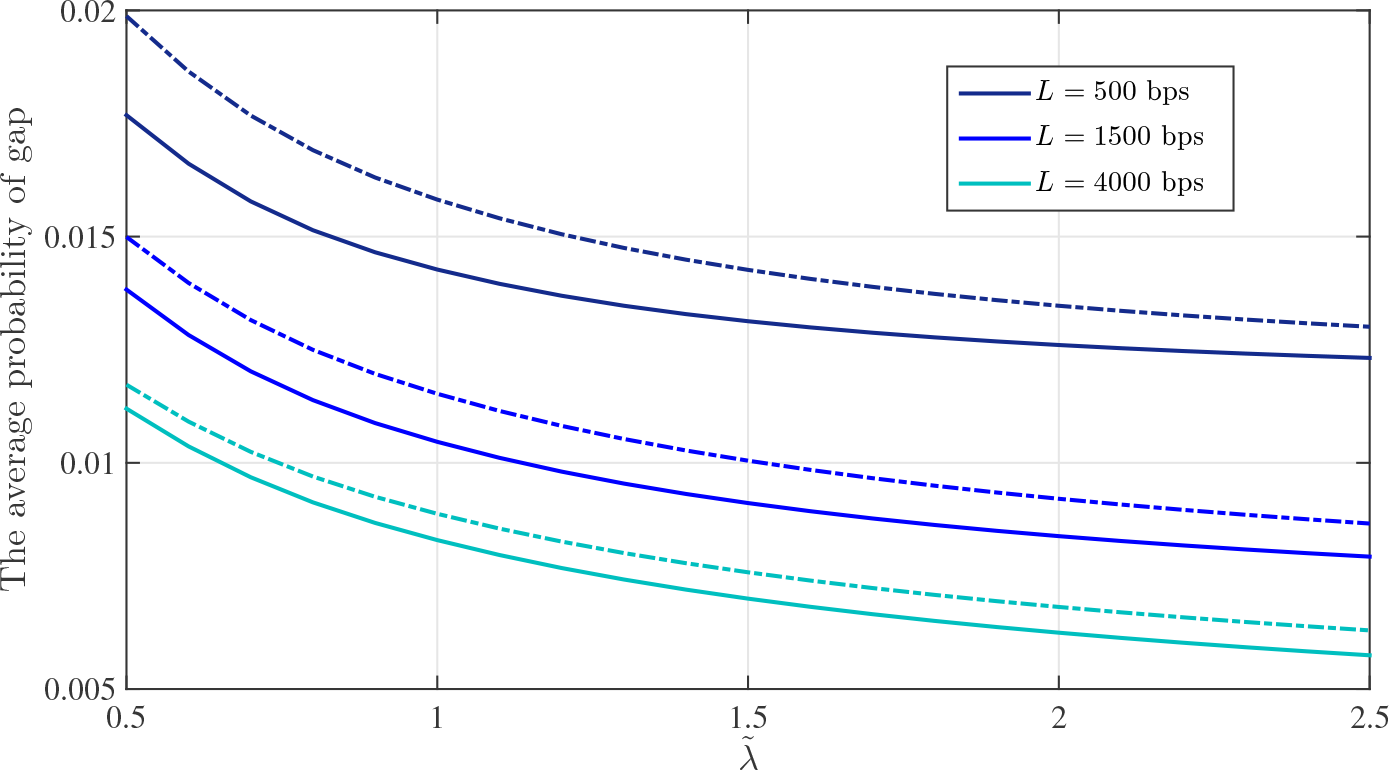
The average probability of gap within Human genome hg19 versus 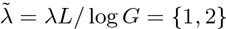 and for different read length *L* = {500,1000, 2000, 3000} bps with p_t_ = 1 and *p_t_* = 0.7 (dotted line).

**Fig. 13.**
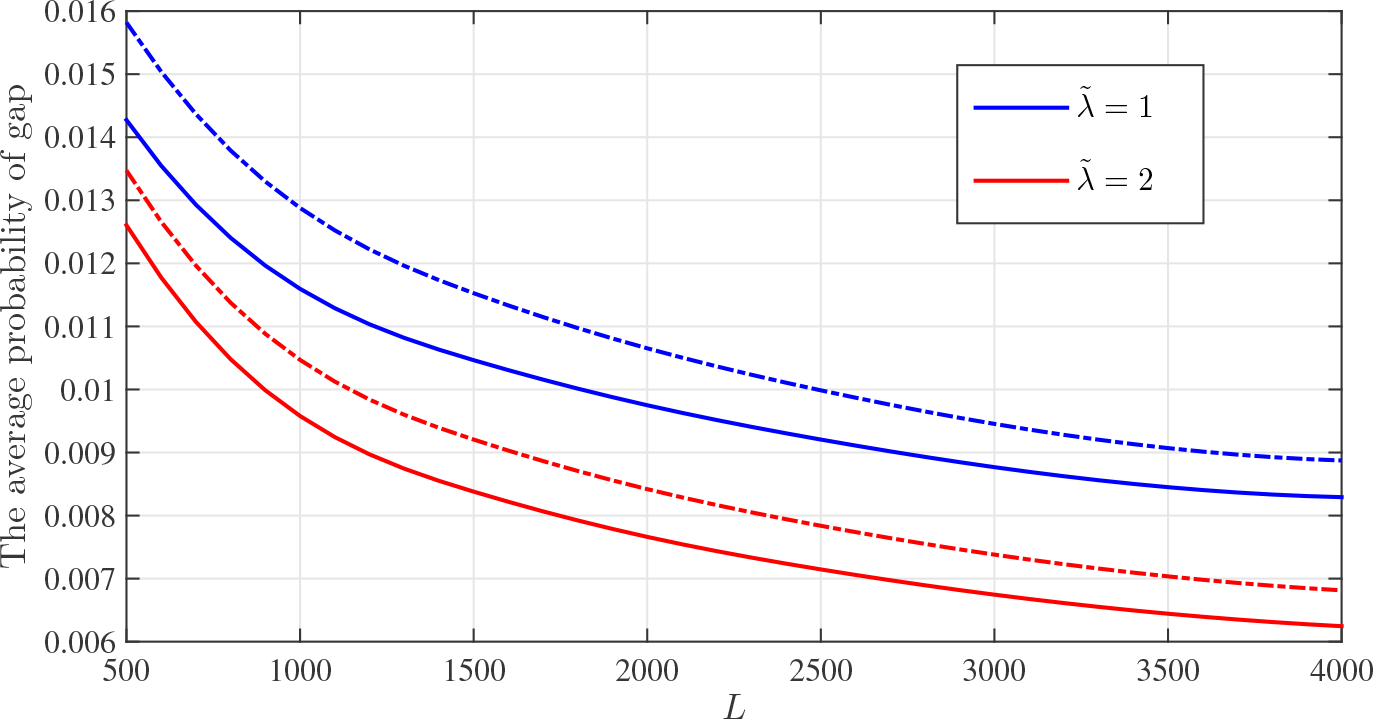
The average probability of gap within Human genome hg19 versus read length *L* and for two values of 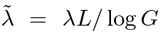 with *p_t_* = 1 and *p_t_* = 0.7 (dotted line).

In each simulation, we also present number of aligned reads by Meta-aligner. Reports of Meta-aligner show its robustness to sequencing errors such that it aligns many reads at its first stage almost correct. We need to read 2*ℓ* bases at the first step of Meta-aligner which increase over-read bases, but most of the mapped reads are located on the genome correctly.

### B. Coverage and Over-read Bases Analysis

Since, number of mapped reads effect on covered bases of the genome, before coverage simulation results we determine number of mapped reads at end of the first stage of Meta-aligner. Figure 14 shows fraction of mapped reads for *ϵ* = {0, 5, 10}%. Results show that most of reads are mapped uniquely to the reference genome.

In Figure 15, the total number of bases not covered by mapped reads (also known as genome gaps) for various sequencing error rates after the first stage of Meta-aligner is presented. By increasing read length more repeat are bridged by reads and the gap fraction is decreased. Also, Figure 16 shows step by step gap faction of the genome for read length *L* = 1000 and different sequencing error rates. This figure shows that using the proposed method, gradually the reference genome is covered by reads. Since, the remaining bases of the genome locate within long repeat regions, they are covered by reads.

**Fig. 14.**
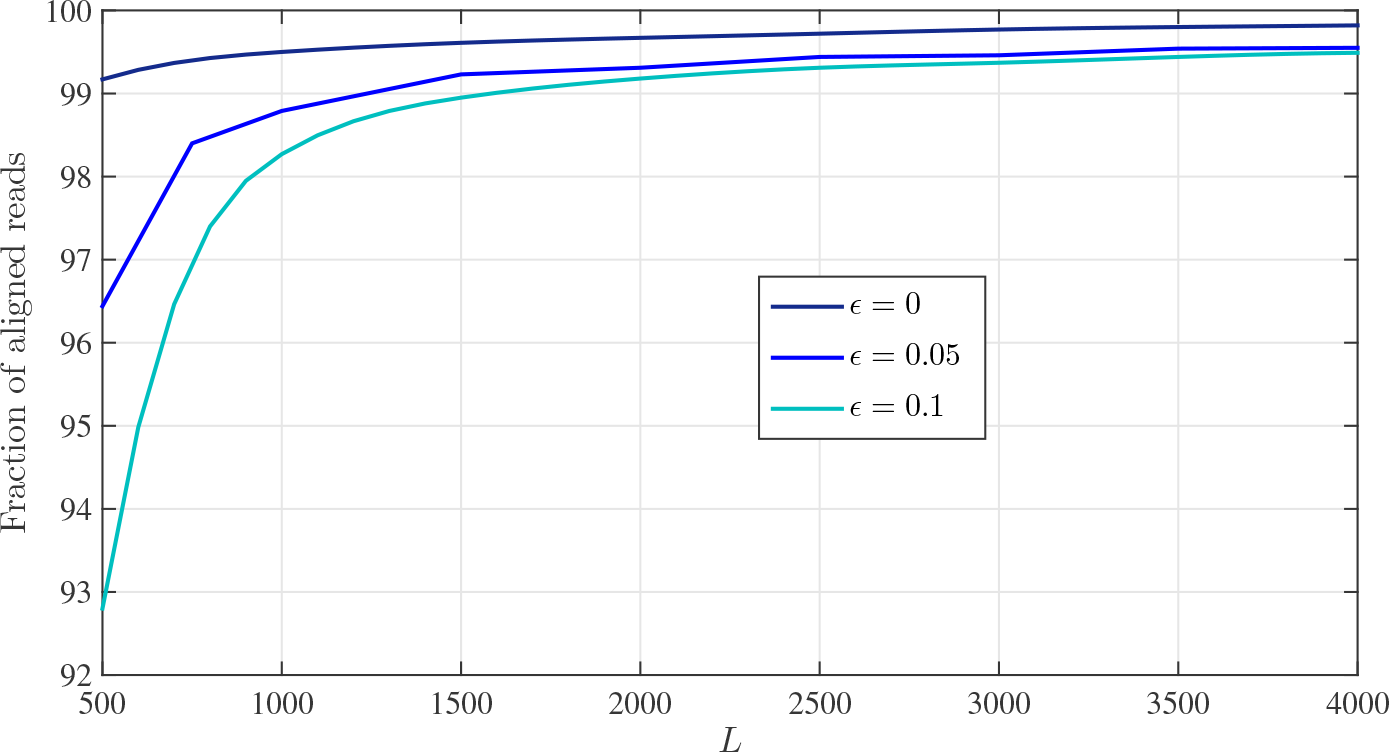
Fraction of mapped reads for different read lengths and sequencing error rates after the first stage of Meta-aligner.

**Fig. 15.**
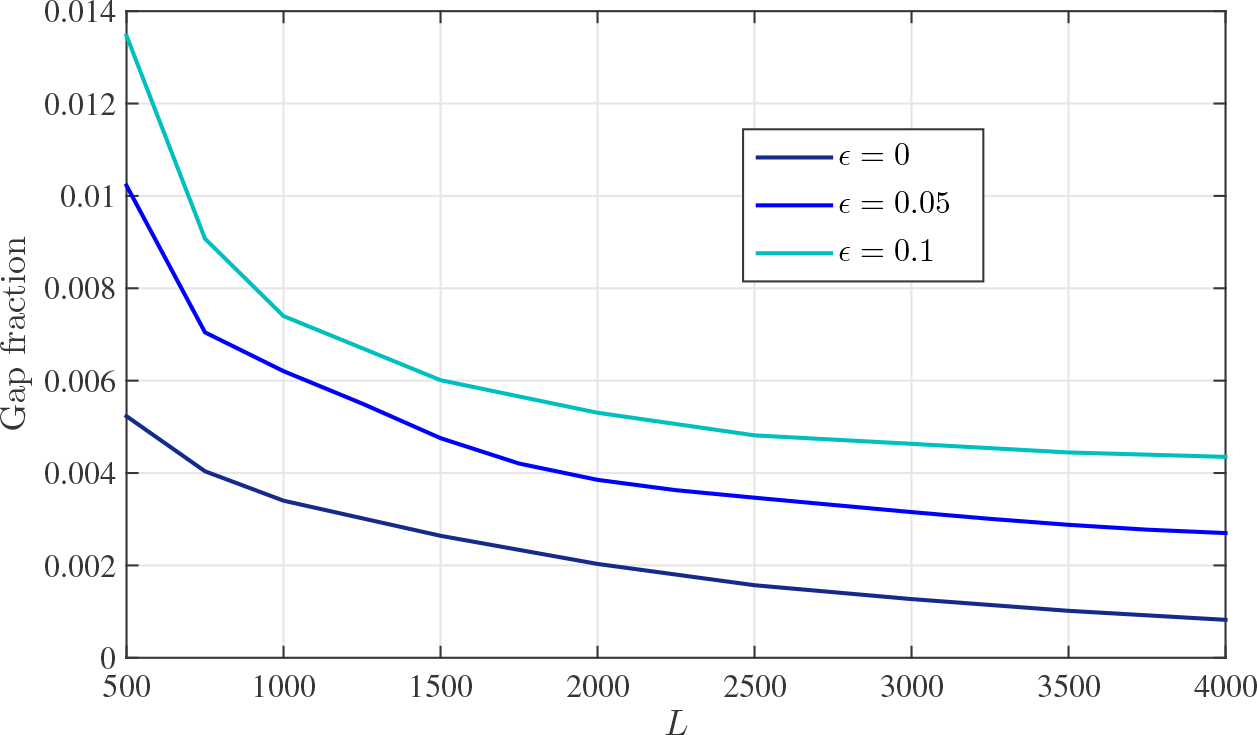
Fraction of gap within the chr19 for different read lengths and different sequencing error rates after the first stage of Meta-aligner.

**Fig. 16.**
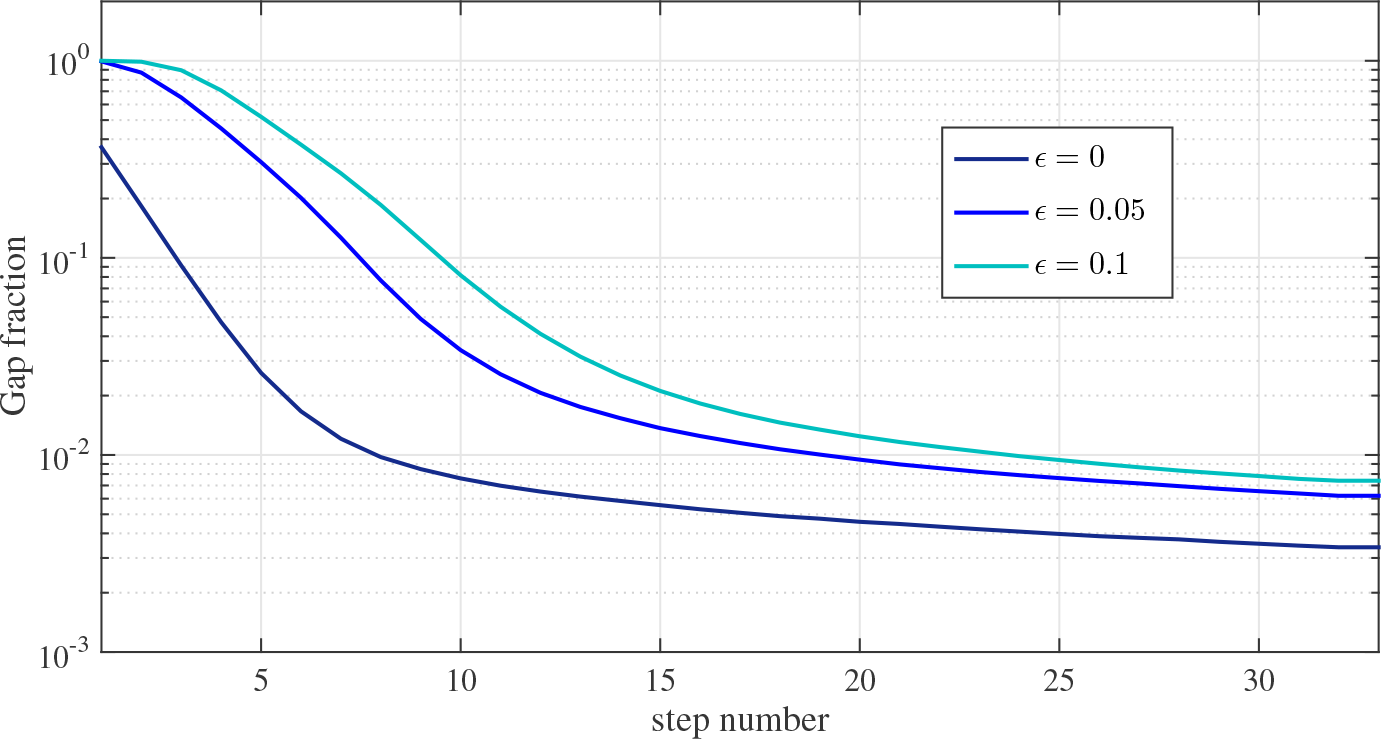
Step-by-step fraction of gap within the chr19 for different sequencing error rates and read length of *L* = 1000.

Figure 17 illustrates the normalized total number of read bases for various sequencing error rates. Note that, for large enough read length number of over-read bases tends to zero and only *O*(*G*) bases needs to be sequenced by sequencer machine. For different values of *ϵ* = {0, 5, 10}%, almost {1.2,6.4,8.7} × *G* bases are read by using Meta-aligner in read length of *L* = 4000 bps, respectively. Also, Figure 18 shows step by step normalized total number of read bases of the genome for read length *L* = 1000 and different sequencing error rates. This figure reveal that after each step some unmapped reads interfere with other mapped reads and total read bases are increased.

**Fig. 17.**
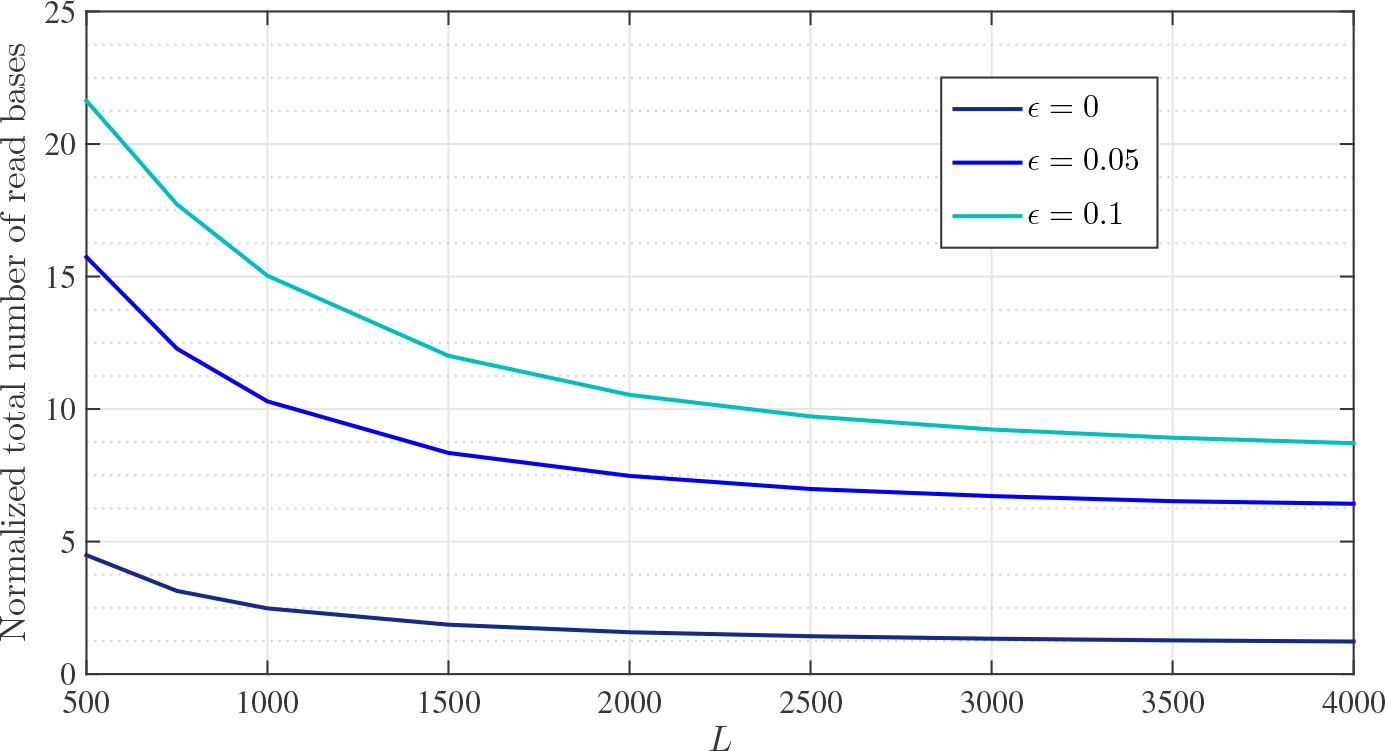
Normalized total number of read bases for various sequencing error rates and read lengths.

**Fig. 18.**
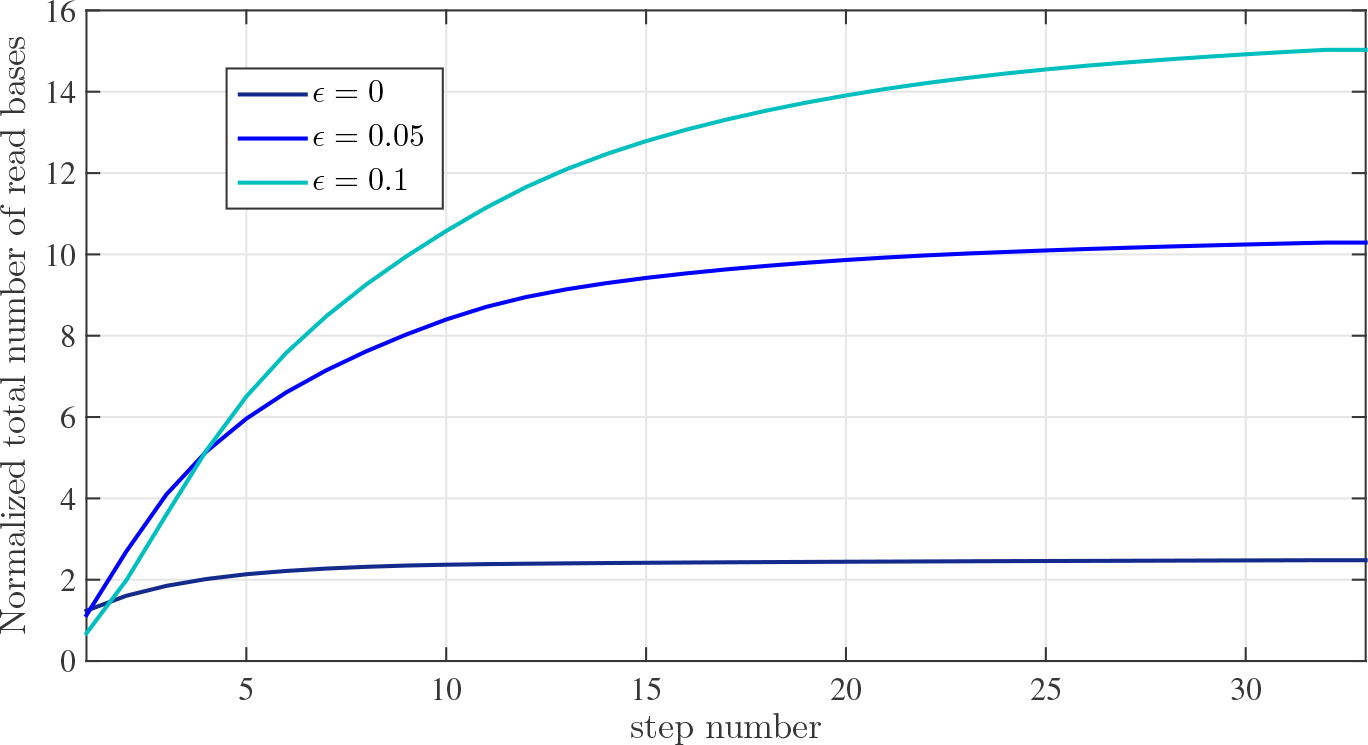
Step-by-step normalized total number of read bases for various sequencing error rates and read length of *L* = 1000.

As these simulation results show, increasing sequencing error rates leads to an increase in the number of over-read bases. This is due to the fact that each read can not be aligned to the genome at shorter lengths and its length is increased iteratively. In addition, it is noticeable that genomes with a larger percentage of repeat patterns naturally lead to a greater level of over-read bases (comparison of chr19 with i.i.d. genome). In such scenarios, less number of reads are uniquely aligned to the genome due to the ambiguity caused by repeating patterns.

## V. CONCLUSION AND FUTURE WORKS

Lander and Waterman have presented the coverage bound based on random sampling of i.i.d. DNA sequence. After sampling, read fragments are sent to the processing part. Under such model, the coverage bound shows that minimum number of reads required for covering the whole genome is *N* ≈ *G* log *G/L*. Equivalently, *N L* ≈ *G* log *G* bases are required to cover the whole genome. In our method, sequencing and processing are combined such that first all fragments are sequenced up to *ℓ* bases, for some *ℓ*, and then the processor maps the fragments that are uniquely mapped to the reference genome. Unmapped reads with non-overlapped reads from the right hand side at the first step are sent back to sequencer for extension to next bases. This procedure is repeated until the process reached the maximum read length *L*. As shown in the paper, through use of such approach, the number of bases read in the sequencing part reduces to *O*(*G*) bases, a reduction by a log *G* factor in comparison with Lander-Waterman coverage bound.

We propose theoretical results for i.i.d. and real genomes with noiseless reads and i.i.d. genome with noisy reads. Also, we have simulated our method for chr19 of Human genome hg19 with different sequencing error rates. Simulation results support the validity of the proposed algorithm and demonstrate our improvement on coverage bound for real genomes.

As future work, we may expand our algorithm to derive more efficient alignment algorithms in terms of complexity and precision. Also, we can extend this method for Denovo sequencing.

